# Autophagic degradation of EIN3 ensures developmental plasticity and recovery from environmental stress in Arabidopsis

**DOI:** 10.1101/2025.08.13.669664

**Authors:** Jeppe Ansbøl, Isolde Riis, Mette Stub, Rim Chaudhury, Emil Otto Kokholm Nielsen, Jonathan Chevalier, Dominique Van Der Straeten, Zhangli Thomsen Zuo, Sjon Hartman, Eleazar Rodriguez

**Author notes:** Now at DTU National Food Institute, Henrik Dams Allé, 202, 2800 Kongens Lyngby. Now at Medpace, Technologielaan 11, 3001 Leuven, Belgium (Jonathan Chevalier).

## Abstract

Ethylene signaling, mediated by the key transcription factor EIN3, regulates diverse developmental processes and stress adaptations, including hypocotyl growth, aging, and submergence tolerance. Autophagy, a cellular recycling process, also facilitates adaptation by reprogramming cellular components. While EIN3 degradation via the proteasome is well established, its connection to autophagy remains unclear. Here, we show that EIN3 turnover is directly regulated by ATG8-mediated autophagy. Consistently, autophagy-deficient plants exhibit impaired EIN3-dependent hypocotyl growth during light-to-dark transitions.

Additionally, EIN3 accumulation contributes to the premature senescence observed in *atg* mutants. Beyond development, our combination of cell imaging, phenotypic analyses, and proteomics reveals that autophagy is essential for EIN3-driven transcriptional reprogramming during submergence. Together, our findings uncover a direct role for autophagy in regulating EIN3 stability, providing mechanistic insight into how this process fine-tunes ethylene responses in growth and stress adaptation.

## Introduction

Cellular adaptation to changing environments depends on the coordinated regulation of macromolecular components, particularly through proteome reprogramming ^1^. Autophagy (*sensu* Macroautophagy), contributes to this process by degrading and recycling intracellular material, thereby supporting dynamic cellular responses to both physiological and stress-related cues ^2^.

Precisely, autophagy has been shown to mediate stress responses like heat stress, salinity, dehydration, submergence, and immunity ^3–8^. Moreover, several reports demonstrate that autophagy is also necessary for proper developmental events like seed germination, lateral root formation, pollen and silique maturation and, aging ^5,9–12^. Autophagy’s participation in these processes is connected to its ability to perform both bulk and selective target degradation via cargo adaptors like NBR1, DSK2, TSPO and EXO70D ^13–16^, enabling the degradative plasticity needed for response optimization. To participate in developmental and stress responses, autophagy must intersect with both endogenous and exogenous signals, allowing for its modulation upon signal perception and enabling it to regulate downstream signaling outputs.

Among these signals, hormones are some of the most dynamic and potent plant regulators, and they have been shown to rapidly enact autophagic activity ^1^. Likewise, several reports indicate that autophagy modulates auxin, cytokinin, ABA and brassinosteroids responses via degradation of key components of these hormone’s signaling pathways(e.g. ^11,13,15,17^).

Another plant hormone shown to activate autophagy is Ethylene (ET, ^1,18^). ET signaling in *Arabidopsis* is complex and has been well described with hormone perception leading to stabilization of the ET master transcription factor ETHYLENE INSESITIVE 3 (EIN3) and its homologues, promoting activation of ET-dependent responses. ET levels influence a range of plant developmental features like skoto-and photomorphogenesis, ripening and senescence ^19–21^. ET also mediates stress responses to flooding, in which gas exchange becomes restricted leading to simultaneous ethylene accumulation and oxygen deprivation (hypoxia) ^22^, having severe consequences for development. Per contra, absence of ET promotes EIN3 ubiquitination and subsequent proteasomal degradation by the SKP1-CULLIN-F-BOX EIN3-BINDING F-BOX 1 (EBF1) and EBF2 ^19,23,24^. Despite the critical importance of EIN3s turnover to control ET-dependent responses, little is known about a putative role for autophagy in this context.

Interestingly, autophagy mutants display phenotypes which intersect with ET signaling, like early senescence ^4^ and hypersensitivity to submergence stress ^3^, which suggests that autophagy might indeed be necessary to modulate ET-dependent responses.

Here we find that autophagy is involved in EIN3 degradation to modulate ET-dependent responses. Our data shows that autophagy is rapidly engaged to promote EIN3 turnover upon perception of different cues like transition from darkness to light or during submergence recovery. Moreover, we show that developmental phenotypes seen in autophagy deficient mutants like altered hypocotyl elongation and premature senescence are caused by misregulation of EIN3. Concurrently, our results suggest that the previously reported sensitivity of *atg* mutants to submergence stress can be explained by overaccumulation of EIN3 and that defective reprogramming of EIN3-dependent submergence responses during recovery contribute to impaired recovery of *atg* mutants after submergence. In sum, we provide novel evidence of autophagic regulation of EIN3 and how it impacts different developmental and stress responses in plants.

## Results

### Autophagy regulates EIN3 levels

To understand if autophagy participates in EIN3 turnover, we verified the steady state levels of EIN3 in *atg2-1* and *atg7-3* mutants by western blot (Figure 1A and Figure S1A). We observed that EIN3 accumulated in these autophagy deficient mutants and importantly, treatment with proteasome inhibitor MG132 led to EIN3 accumulation, the latter being in agreement with previous findings detailing proteasomal turnover of EIN3^23,24^. These data indicate that EIN3 turnover is regulated by both degradation pathways. To ensure that EIN3’s accumulation in *atg* mutants is not caused by transcriptional upregulation, we quantified *EIN3* transcript levels by RTqPCR and observed no statistical difference from Col-0 (Figure 1 B; Figure S1B). To further support the role of autophagy in EIN3 turnover, we crossed a line expressing EIN3-GFP fusion protein driven by its native promoter (*pEIN3::EIN3-GFP*) with *atg2-1* and quantified the GFP fluorescence levels of those plants and the parental line. In agreement with our western blot data, fluorescence quantification revealed that EIN3-GFP accumulates in *atg 2-1* when compared to the parental line (Figure 1 C, D). Together, these results support a role for autophagy in the modulation of EIN3 levels.

**Figure 1:**
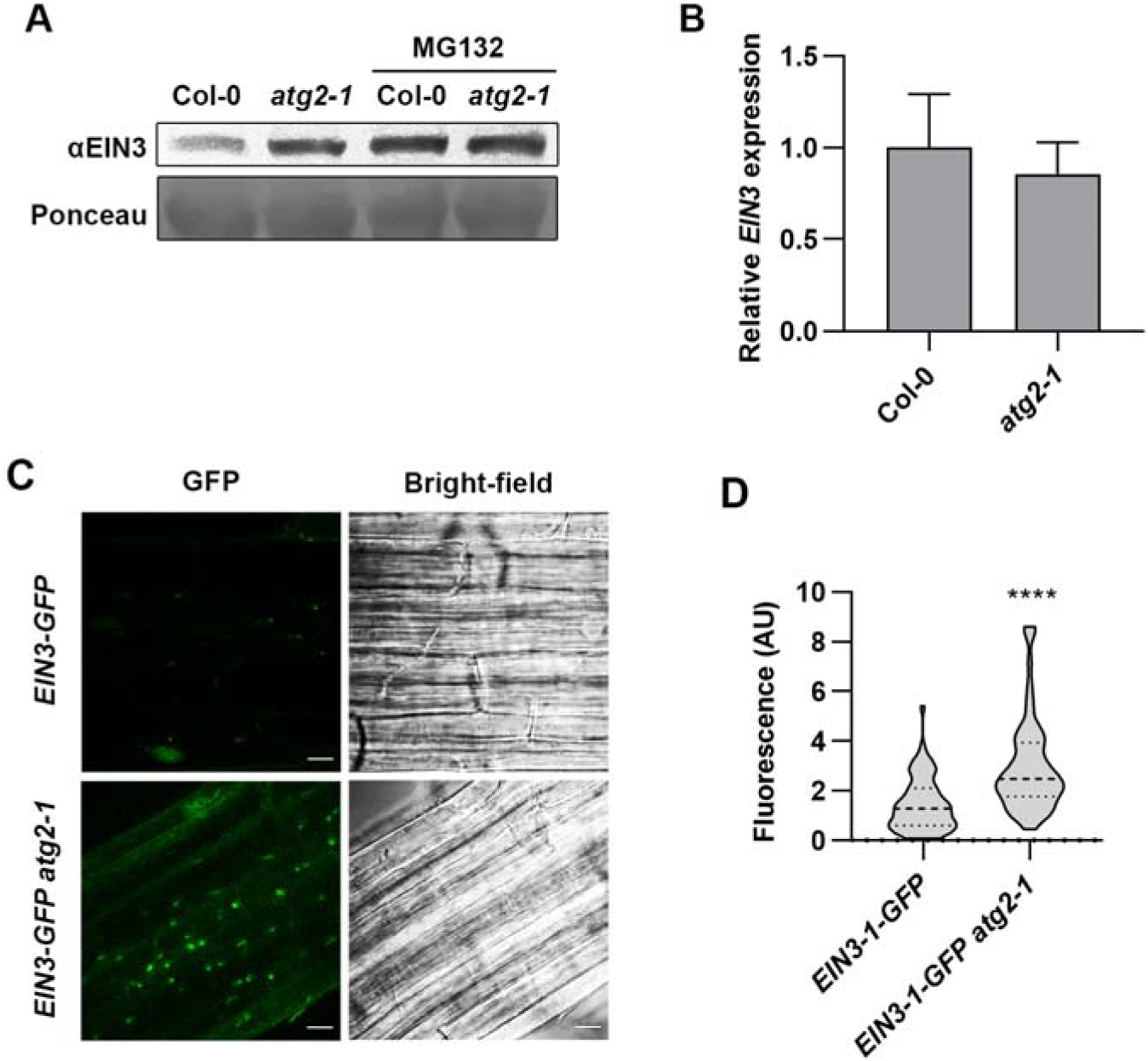
EIN3 accumulates in autophagy deficient *atg2-1* mutants. **A)** Immunoblots from proteins of Col-0 and *atg2-1* seedlings treated with MG132. The experiment was repeated twice with similar results. **B)** Relative expression of *EIN3* in *atg2-1* and relative to Col-0. *Actin2* was used to normalize the expression. **C, D)** Fluorescence quantification for *EIN3-GFP* and *EIN3-GFP atg2-1.* At least 6 plants were analyzed per condition.

### ATG8 interacts with EIN3

Having established that EIN3 accumulates in *atg* mutants and is likely targeted to degradation by this pathway, we wanted to confirm this by performing protein-protein interaction studies. Using Arabidopsis stable lines expressing GFP-ATG8a, we performed co-immunoprecipitation (Co-IP) and probed for native EIN3. As seen in Figure 2 A, there is an enrichment of EIN3 when GFP-ATG8a is immunoprecipitated, but not in the negative control. In support of our Co-IP data; Bimolecular fluorescence complementation (BiFC) assays showed reconstitution of fluorescent signal between nCFP-ATG8a and EIN3-cVenus in both the nuclei and cytoplasm but not when EIN3-cVenus was co-expressed with nCFP-GUS(Figure 2 B).

**Figure 2:**
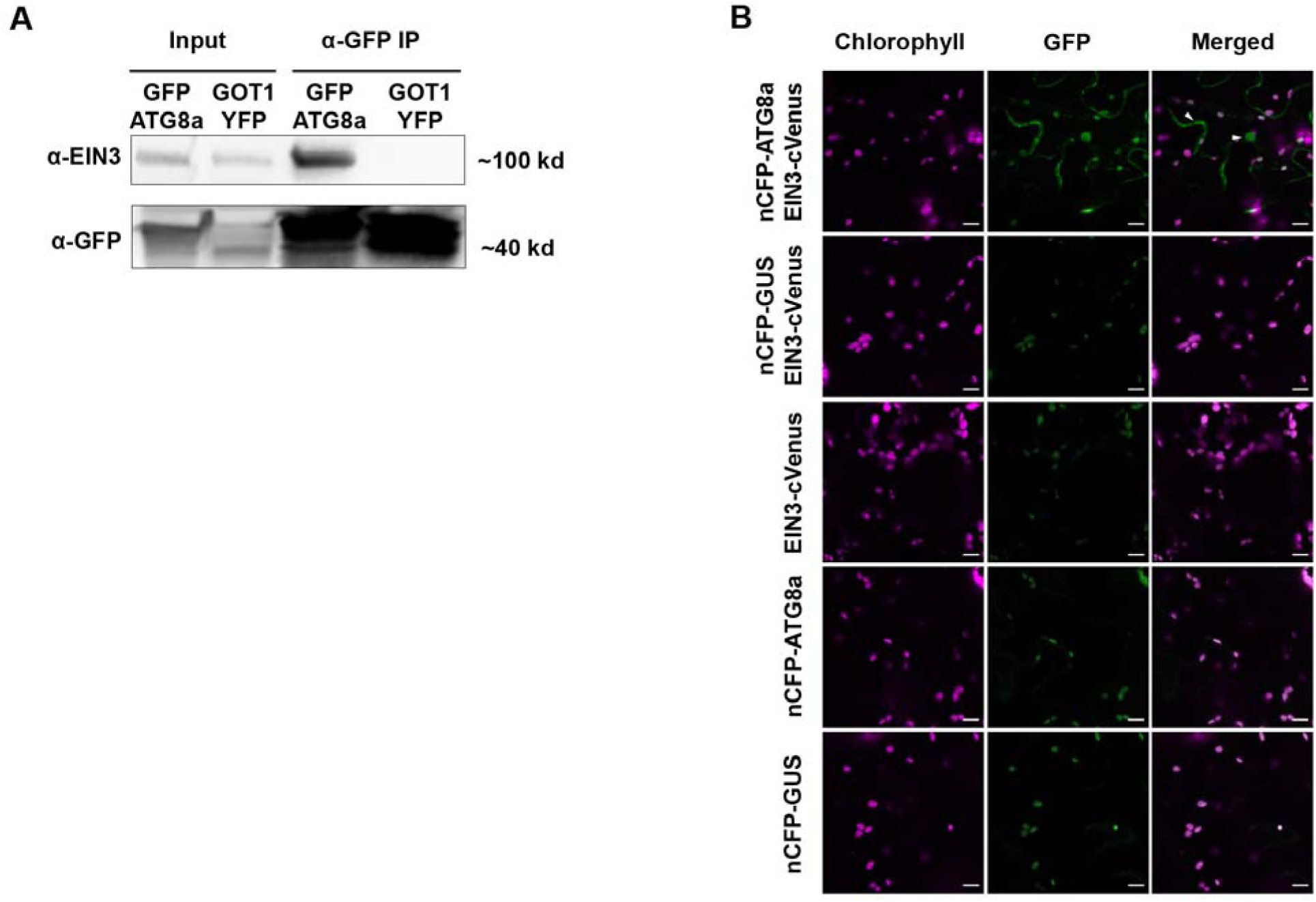
Autophagy protein ATG8 interacts with EIN3. **A)** ATG8a and EIN3 association tested by Co-IP assay. Protein extracts from stable *Arabidopsis* lines expressing GFP-ATG8a or GOT1-YFP were immunoprecipitated with anti-GFP antibody. The membranes were then probed for either anti-EIN3 or anti-GFP antibodies. **B** Bimolecular fluorescence complementation experiments showed oligomerization of EIN3 and ATG8a in *N. benthamiana.* White arrows indicate signal reconstitution. Images were taken 48 hpi. cCFP-GUS was used as control. Scale bar: 10 μm.

### Dynamic conditions promote EIN3 localization in autophagosomes

As our data indicate that autophagy promotes EIN3 turnover, we wanted to test conditions leading to EIN3 colocalization with ATG8 puncta. To this end, we crossed stable lines expressing EIN3-GFP with mCherry-ATG8 and performed confocal microscopy on different scenarios (Figure 3). Under standard conditions, we observed a modest colocalization of EIN3-GFP and GFP-ATG8 colocalization, which might relate to the low signal of EIN3-GFP (Figure 2 A). As the ET precursor 1-aminocyclopropane-1-carboxylic (ACC) is known to stabilize EIN3 ^25^ we treated seedlings with ACC. As expected, ACC treatment led to enhanced EIN3 signal but interestingly, not to an increase in colocalization with GFP-ATG8a (Figure 2 B). We then reasoned that replacing ACC containing media for hormone free media could trigger colocalization, and precisely in that set up we observed significant increase in colocalization between the 2 proteins, indicating that upon ACC removal, EIN3 is targeted to autophagic puncta (Figure 2 C). As with ACC treatment, dark growth conditions are known to promote EIN3 stabilization; this can be rapidly reverted by switching to light growth conditions, (Hui Shi et al. 2016). In agreement with those findings, we saw an increase in colocalization of EIN3-GFP with mCherry-ATG8a upon switching seedlings from dark to light growth conditions (Figure 2 D). It should be noted that given the fast turnover of EIN3 under light growth conditions, we had to add vacuolar protease inhibitors (E-64d and Pepstatin A) right before switching the plants from dark to light growth conditions and imaging, in order to detect colocalization of EIN3-GFP and mCherry-ATG8. Importantly, light grown seedlings pretreated with vacuolar protease inhibitors before imaging did not display enhanced colocalization of the 2 proteins (Figure 2 E). This indicates that the transition from dark to light growth conditions triggers rapid localization of EIN3-GFP to ATG8a puncta (Figure 3). In addition, we tested whether submergence and recovery from submergence stress also affected colocalization of EIN3-GFP with mCherry-ATG8a, as these are known to require tight regulation of ET-dependent responses ^21,26–28^.

**Figure 3:**
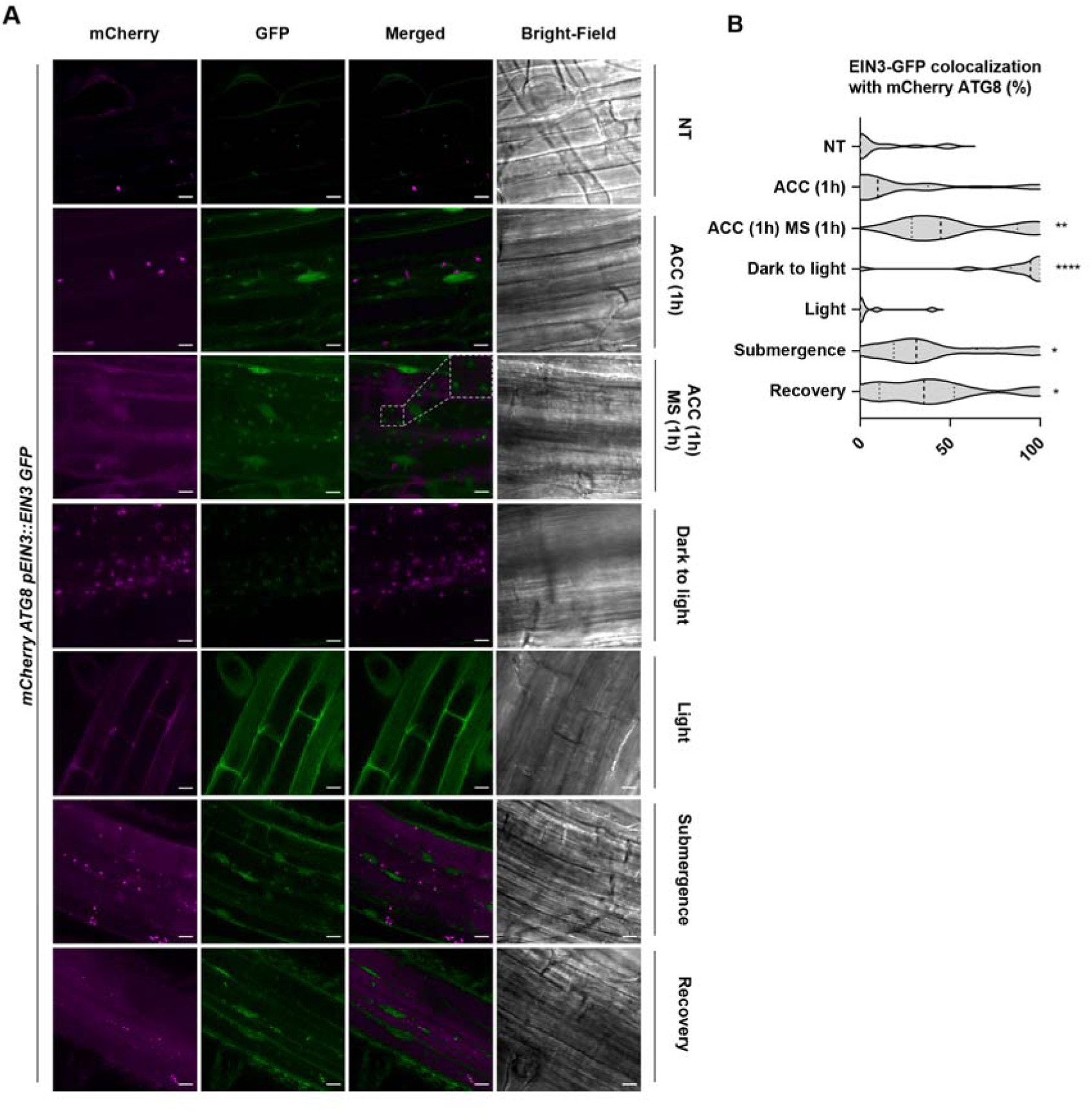
Confocal microscopy showing EIN3-GFP colocalizes with mCherry-ATG8 during different stresses. **A)** Confocal microscopy of EIN3-GFP and mCherry-ATG8 under various treatments. **B)** Violin plot of colocalization in % of EIN3-GFP to mCherry-ATG8 foci. Asterisks denote Dunn’s multiple comparisons test (*0.05; **0.01, ****0.001). The treatments are MS (NT); 10 µM ACC for 1 hour (ACC 1h), 10 µM ACC for 1 hour and subsequently replaced with MS for 1 hour (ACC1h MS1h); seedlings grown at standard conditions before the lights are switched on and then treated with vacuolar protease inhibitor 10 μM E-64d and 1 μM pepstatin A for immediate imaging (Dark to light); light grown seedlings treated with 10 μM E-64d and 1 μM pepstatin A for immediate imaging (Light); overnight submergence with no light (Submergence) or 1 hour after removal from dark submergence (Recovery). At least 4 seedlings were analyzed per condition. Scale bar = 10 µm

Interestingly, both conditions displayed similar levels of colocalization, which were significantly higher than in not treated plants (Figure 2 F, G). Collectively, our microscopic observations for colocalization between EIN3-GFP and mcherryATG8a, support autophagy-mediated EIN3 turnover under dynamic growth conditions.

### Hypocotyl growth in darkness of *atg* mutants are EIN3-dependent

Given the role of autophagy in EIN3 degradation, we wanted to check if there were phenotypical consequences for EIN3 misregulation in *atg* mutants. To test this, we examined EIN3-dependent developmental phenotypes in *atg* mutants. A well characterized EIN3-regulated phenotype is skotomorphogenesis, in which dark grown seedlings elongate their hypocotyl until light is perceived, causing a shift towards photomorphogenesis ^29^. Because the switch from skoto-to photomorphogenesis induces rapid EIN3 turnover ^21^, we wondered if hypocotyl elongation could be defective in *atg* mutants. To investigate this, we grew Col-0*, atg2-1, ein3-1* and *atg2-1 ein3-1* seeds and tracked hypocotyl elongation in darkness for 2.5 days, followed by transition to light regime, using an automated plant imager SPIRO ^30^. Our data shows that *atg2-1* exhibited both a reduced growth rate and shorter hypocotyls compared to Col-0 and *ein3-1*. Notably, this growth defect was rescued in the *atg2-1 ein3-1* double mutant (Figure 4 D, E). This phenotype is not due to delayed germination, as all genotypes showed comparable growth during the first 30 hours (Figure S2). Beyond this period, *atg2-1* seedlings displayed a markedly slower rate of hypocotyl elongation (between 20 and 40h), with a slope 2 times lower than those of the other genotypes (0.25 for *atg2* vs 0.5 for Col-0). Transition from dark to light growth regime should lead to rapid halt of hypocotyl elongation ^31^, which is regulated by EIN3. To check if autophagy impacts the switch between skoto-and photomorphogenesis, we calculated the slope of the growth curve for the first 50 minutes after light switched on for all the genotypes. As expected, we observed a deceleration of hypocotyl growth in all genotypes once the lights were switched on; however, the deceleration was at least 1.6 times slower in the *atg2* single mutant (slope = −0.014) compared to Col-0 (slope = −0.022), *ein3-1* (slope = −0.027), and the *atg2 ein3-1* double mutant (slope = −0.025). As the *atg2-1 ein3-1* double mutant displayed WT-like response, these results suggest that autophagy is needed to optimize hypocotyl elongation during dark-to-light transition, likely through regulation of EIN3. Strengthening this proposition, a western blot analysis during the transition between skoto-to photomorphogenesis showed that while Col-0 efficiently degraded EIN3 upon light exposure, this protein persisted up to in *atg* mutants (Figure 4 A, Figure S3).

**Figure 4:**
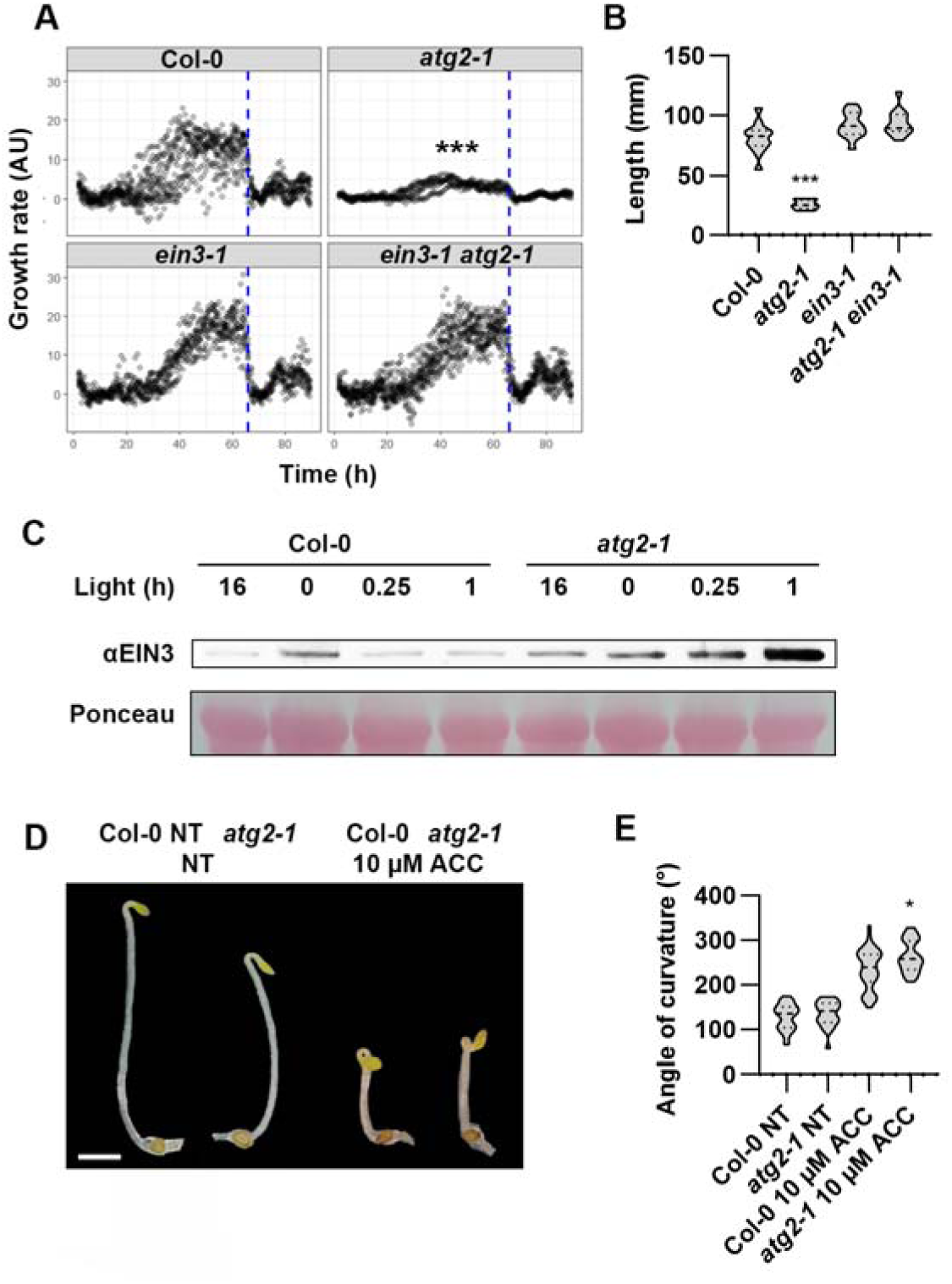
Hypocotyl growth in darkness is *atg2-1* dependent. **A)** Growth rate of Col-0, *atg2-1, ein3-1 and atg2-1 ein3-1* and their response to light. The blue striped line indicates the beginning of light exposure. Asterisks depict statistical significance from Col-0 according to a linear mixed model (***0.001). At least 13 plants were analyzed. **B)** Hypocotyl length from (A). Asterisks depict statistical significance from Col-0 according to Dunn’s multiple comparisons test (***0.001) **C)** Immunoblot of Col-0 and *atg2-1* seedlings for EIN3 at different hours of light exposure. Experiment has been replicated once with similar results. **D**) Triple response in Col-0 and *atg2-1*, with the angle being the apical hook angle of the hook. Scale bar = 1 cm **E)** Curvature of the hook from (D). The asterisks depict statistical significance from Col-0 according to a t-test (*0.05). At least 21 plants were analyzed.

Encouraged by these results, we next tested another well-known EIN3 dependent response, ET-induced hypocotyl hook formation ^32^. As such we measured the hypocotyl hook angle of Col-0 of *atg2-1* seedlings grown with or without supplementation with ACC (Figure 4 D, E).Under untreated conditions the 2 genotypes were statistically undistinguishable (P>0.05), ACC treatment on the other hand led to exaggerated hook formation in *atg2-1* as compared to Col-0 (Figure 4 D, E), suggesting increased ET-sensitivity in *atg2-1*. In summary, the data presented in figure 4 indicate that autophagy plays an important role during ET-dependent early developmental events via modulation of EIN3 levels.

### Early senescence in *atg* mutants is EIN3 dependent

A common phenotype for both EIN3 accumulation and autophagy deficiency is early senescence ^4,20^, and given our data, we wondered if the early senescence of *atg* mutants was caused by EIN3 accumulation. To address this, we monitored the progression of development through to the onset of senescence across key genotypes, including wild-type, autophagy-deficient, and EIN3-related lines (Figure 5). As previously reported, *atg* mutants display early senescence at 4-5 weeks under l6-8 photoperiod ^4,5^, however *atg5 x ein2* double mutants were described as still displaying early senescence, ruling out ET as a cause for the early senescence phenotype. In contrast to those findings, our results show that early senescence of *atg* mutants was EIN3-dependent as neither of the *atg ein3-1* double mutants displayed the characteristic early senescence (Figure 5 A, B, Figure S4) To support this finding, we measured the levels of the senescence marker senescence associated gene 12 (SAG12) ^33^ in the same plants as above by western blot. Consistent with the developmental phenotype, SAG12 accumulated in *atg2-1* mutants in an EIN3-dependent manner (Figure 5 C). Furthermore, as an indicator of senescence we measured the chlorophyll levels of rosette leaves in 5-week-old plants and found a significant decrease in chlorophyll A and B content in *atg2-1* compared to Col-0, *ein3-1* and *atg2-1 ein3-1* (Figure 5 D), again indicating that the premature aging of *atg* mutants is caused by EIN3 misregulation.

**Figure 5:**
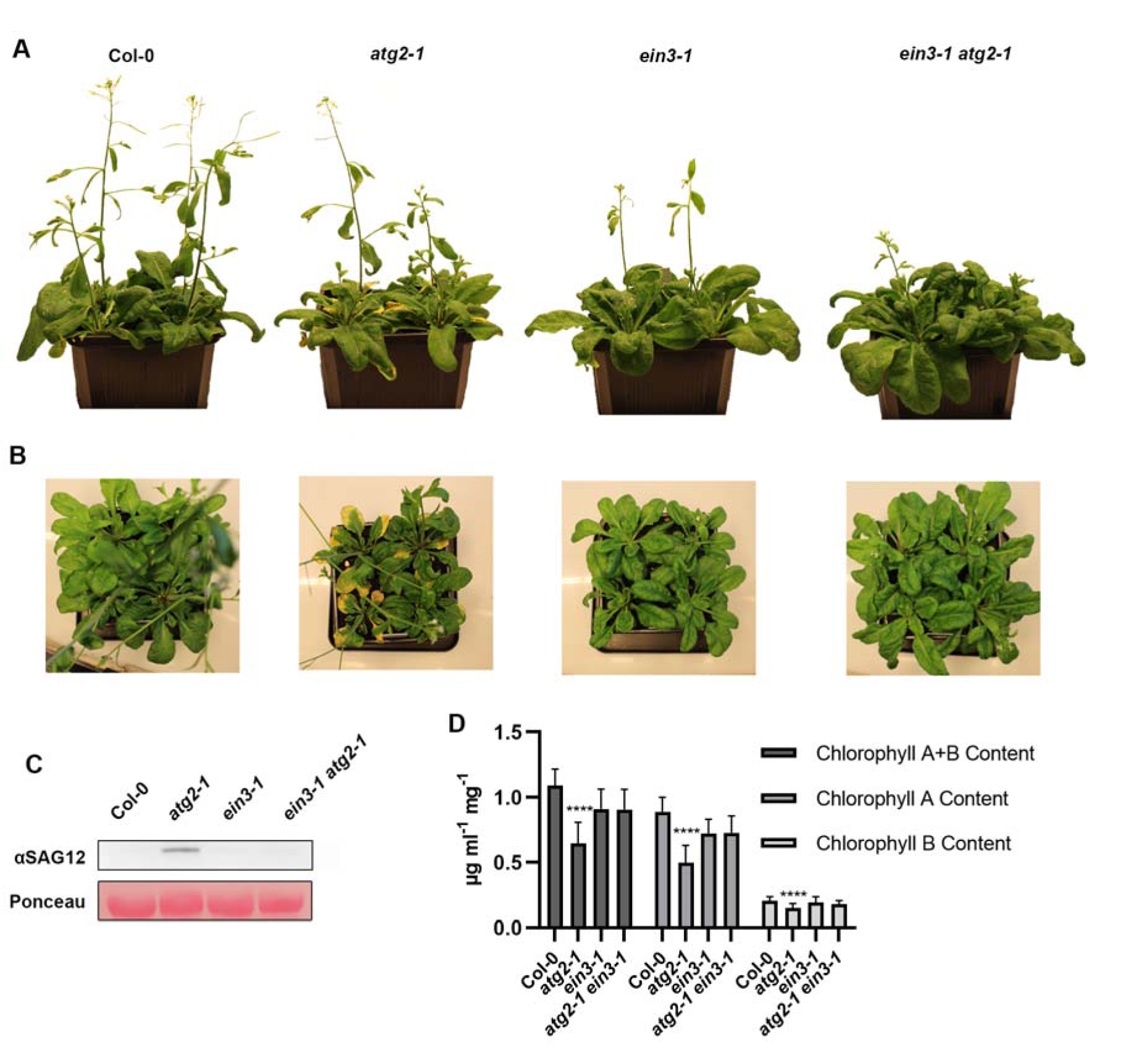
Early senescence of *atg* mutants is EIN3 dependent. **A, B)** 5-week-old *Arabidopsis* with different genotypes grown at standard conditions. **C)** Immunoblot of 5-week-old *Arabidopsis* was probed for SAG12. The experiment was repeated 3 times with similar results **D)** Chlorophyll A, B, and total (A+B) content (µg mllJ¹ extract mglJ¹ fresh weight) (A). Values are from a mean of 10 different plant rosette leaves. Asterisks depict statistical significance from Col-0 according to a Tukey’s multiple comparison’s test (***0.001).

### Autophagy mutants’ sensitivity to submergence is EIN3 dependent

Besides developmental features, ET is also a major regulator of diverse stress response, such as submergence ^34^. Because autophagy is important in eukaryotes for hypoxic responses ^35,36^ and as *Arabidopsis* autophagy mutants have been previously shown to be sensitive to submergence ^3^, we decided to test if these phenotypes were caused by EIN3 misregulation. To verify this, we submerged Col-0, *atg-* and *ein3* related genotypes for one day in darkness and then let them recover under standard growth conditions. As expected, *atg* single mutants had a visible senescing phenotype and had less chlorophyll content than *ein3-1*, *ein3-1 atg* double mutants and Col-0 after submergence. Importantly, the *atg ein3-1* double mutants showed a rescue of the senescence and chlorophyll deficiency observed in *atg* single *mutants* (Figure 5 A, B; Figure S5). This suggests that the submergence phenotype in *atg* mutants is likely linked to its role in the regulation of EIN3.

Because the sensitivity of *atg* mutants to submergence stress was exacerbated during the recovery phase, we wondered if a defect in termination of EIN3-dependent response in the *atg* mutants could explain this sensitivity. To determine how autophagy might play a role in reprogramming the proteome during submergence and recovery we looked at the hypoxia marker alcohol dehydrogenase (ADH) and sucrose synthase 1 (SUS1) which are upregulated during submergence ^37^ and ADH is known to be down-regulated transcriptionally during recovery ^38^. As seen in figure (Figure 6 C), ADH was accumulated in Col-0 and *atg2* immediately after submergence, consistent with is known role in submergence responses. Interestingly, ADH was barely detectable in *ein3-1*-related genotypes, suggesting that EIN3 is necessary for ADH accumulation during submergence. After 3 days of recovery, the levels of ADH were undetectable in Col-0 or *ein3-1* related phenotypes, but strikingly, it had further accumulated in *atg2-1*. Contrary to ADH, SUS1 levels seemed to decrease during submergence and accumulate during recovery for both Col-0 and *atg2-1*. Genotypes related to *ein3-1* displayed lower levels of SUS1 regardless of the condition tested, suggesting that as for ADH, the expression of this protein is EIN3-dependent. Importantly, during recovery SUS1 levels in *atg2-1* were 3 times higher than for Col-0, indicating that autophagy is needed to modulate the protein levels under recovery and together with the ADH data, suggesting that autophagy is important for correct modulation of the proteome during submergence and recovery. As our data also shows enhanced co-localization of EIN3 to ATG8 during and after submergence (Figure 3) and as EIN3 levels are downregulated during recovery in an autophagy-dependent manner (Figure 6D), altogether these results suggest that autophagy has an important role during the recovery period after submergence; namely to ensure that EIN3-dependent submergence responses are terminated upon reoxygenation.

**Figure 6:**
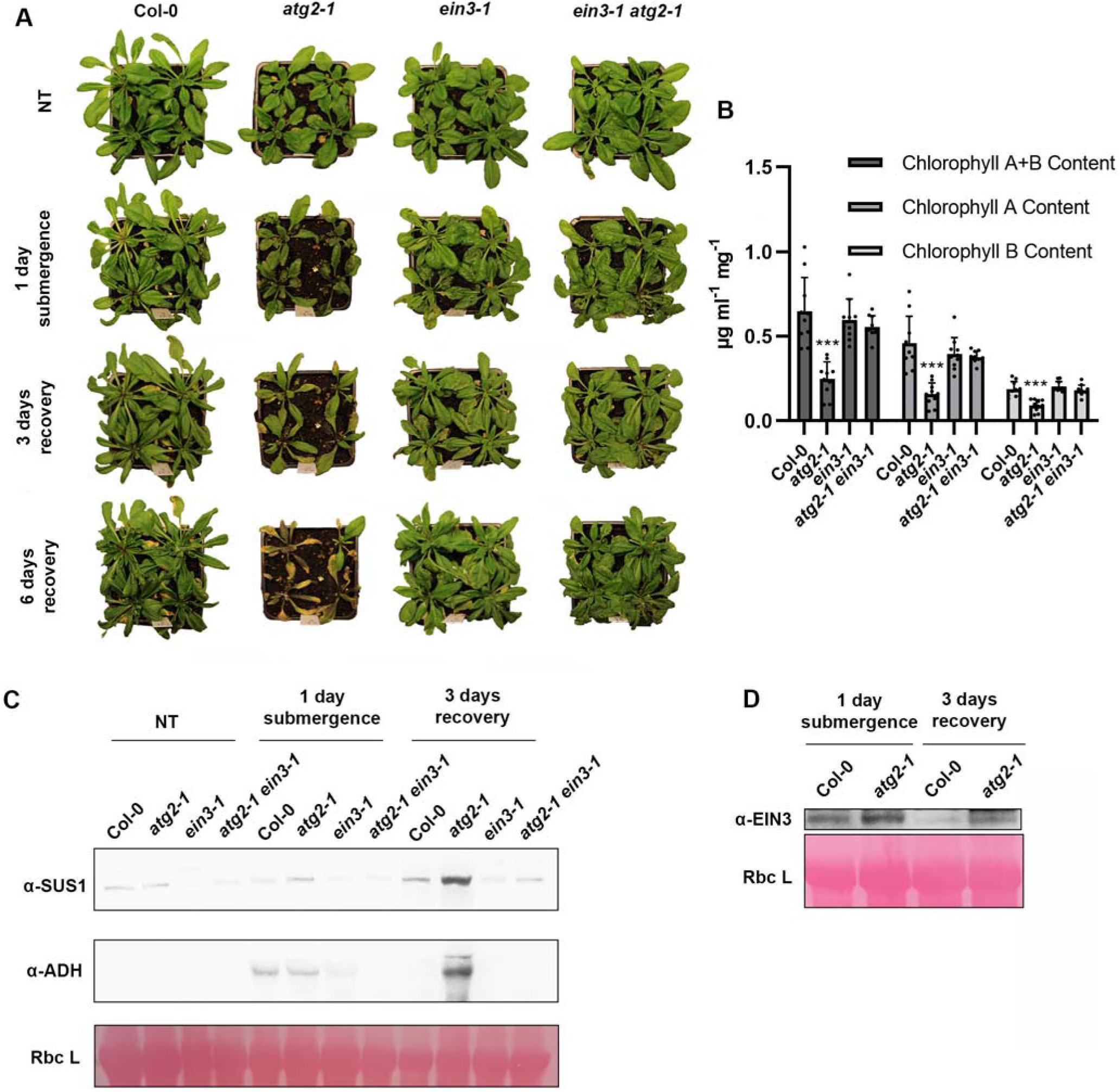
Submergence stress of *atg2-1* is EIN3 dependent. **A)** 4-week-old *Arabidopsis* before submergence, (NT), immediately after 1 day of submergence and 3-or 6-days recovery, **B)** Chlorophyll A, B, and total (A+B) content (µg mLL¹ extract mgL¹ fresh weight) after 6 days of recovery. Values are from a mean of 10 different plant rosette leaves. **C)** Western blots from Col-0, *atg2-1* (a2), *ein3-1* and *atg2-1 ein3-1* before submergence (NT), immediately after 1 day of submergence and 3-days recovery from the submergence. The blot was probed with anti-SUS1 and anti-ADH native antibodies. RuBisCO large subunit (Rbc L) as stained with Ponceau S was used as loading control. **D)** Western blots from Col-0 and *atg2-1* plants exposed to 1 submergence and after 3-day recovery periods. The blot was probed with anti-EIN3 native antibody. RuBisCO large subunit (Rbc L) as stained with Ponceau S was used as loading control. Western blots in **C&D** were repeated 2 with similar results.

### Autophagy modulates EIN3-dependent proteome remodeling during submergence and recovery mutants

Given that our results indicated a requirement for autophagy in proteome remodeling during submergence and recovery, we sought a more detailed view of these dynamics. To this end, we performed label-free mass spectrometry on whole-protein extracts from Col-0, *atg2-1*, and *ein3*-related genotypes under control, submergence, and 3-day recovery conditions. As expected, proteomic analysis revealed pronounced differences in *atg2-1* compared to Col-0, *ein3-1*, and *atg2-1 ein3-1* after submergence, with magnification of these difference following recovery (Figure 7, Figure S6). Unsupervised clustering of differentially abundant proteins identified a group of eight clusters sharing a broadly similar profile: submergence-induced accumulation that was at least partially EIN3-dependent, followed by downregulation during recovery in Col-0 and *ein3-*related genotypes, but persistent or elevated levels in *atg2-1*. These included proteins involved in cell wall remodeling —such as multiple XYLOGLUCAN ENDOTRANSGLUCOSYLASE/HYDROLASE (XTH18, XTH19, XTH24, XTR6), β-GALACTOSIDASE (BGAL4),—as well as defense, senescence and stress markers like ENHANCED DEFENSE SUSCEPTIBILITY (EDS16), PHYTOALEXIN DEFICIENT (PAD3), and ARABIDOPSIS NDR1/HIN1-LIKE 10; YELLOW-LEAF-SPECIFIC GENE 9; (NHL10/YSL9), LIPOXYGENSASE (LOX1), GLUTATHIONE S TRANSFERASE (GSTU4), HEMOGLOBIN (HB1)^39,40^ and ADH (which we had confirmed earlier, Figure 6C) (Figure S7A). This pattern indicates that *atg2-1* retains a stress-associated proteome during recovery, consistent with the visible senescence phenotype observed in these plants (Figure 6A).

**Figure 7:**
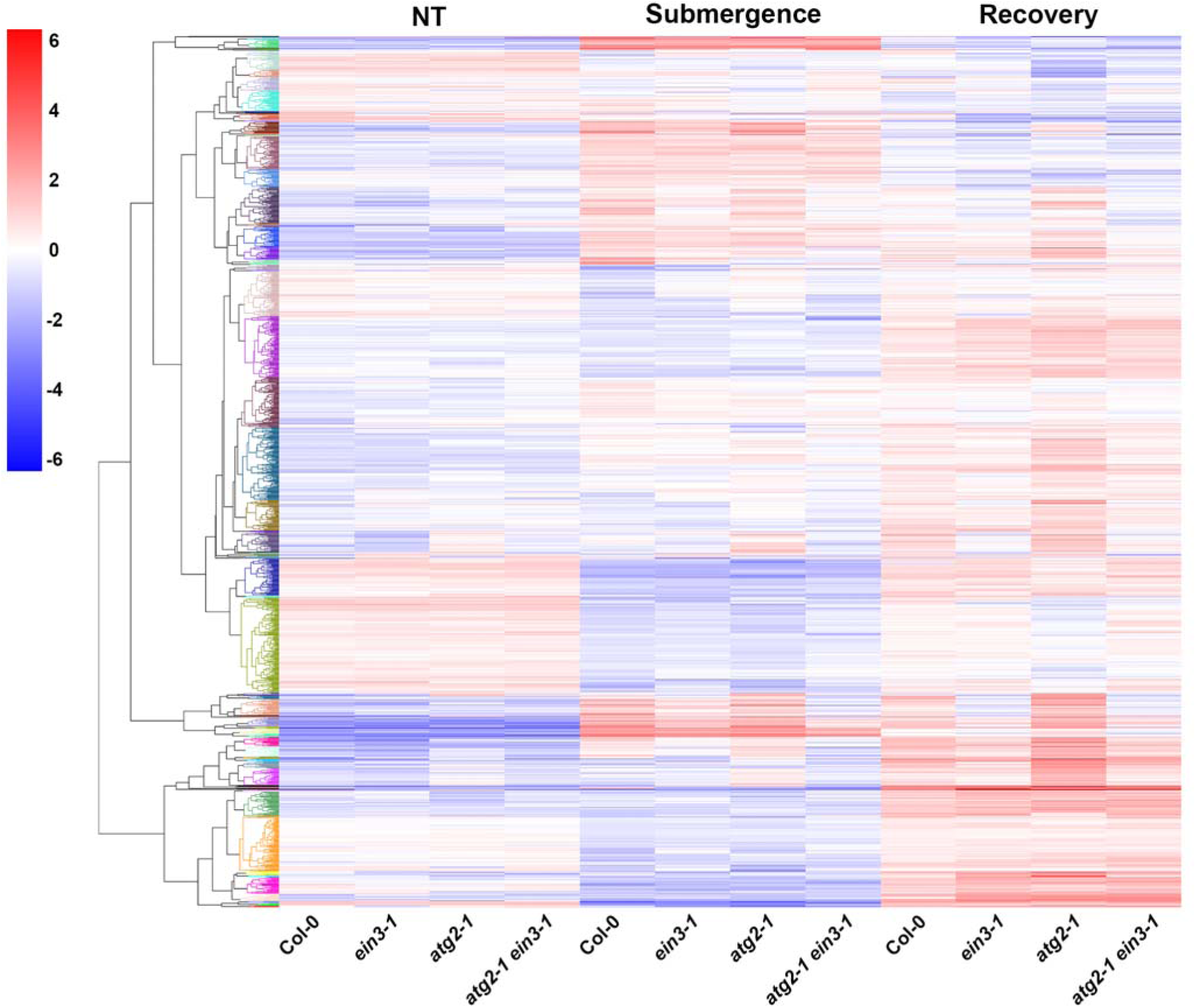
Heatmap of gene expression in 4-week-old plants subjected to submergence and recovery treatments. Plants were submerged in darkness for 1 day (Submergence) and then returned to standard growth conditions for 3 days (Recovery). Untreated (NT) controls were maintained under standard growth conditions for the full 4-day period. Values represent the average of 4 biological replicates per condition and genotype. Hierarchical clustering was performed on proteins that were significant in at least one ANOVA or unpaired t-test comparison. In each clustermap, selected features were mean-normalized.

Importantly, these proteins do not accumulate in the *atg2-1 ein3-1* double mutant, suggesting that autophagy is required to restrain EIN3-dependent stress signaling after submergence, thereby facilitating the proteomic shift toward recovery.

Conversely, a different subset of proteins was downregulated during submergence across all genotypes, followed by robust upregulation during recovery in all genotypes but *atg2-1*, which showed either mild or no accumulation of these proteins. These clusters include numerous proteins involved in cell wall biosynthesis like: CELULLOSE SYNTHASE (CESA1/4/5/6), β-GALACTOSIDASE (BGAL3), WALL ASSOCIATED KINASE (WAK1/2), development: BRI1 EMS SUPRESSOR (BES1), ELONGATED HYPOCOTYL (HY5), KNOTTED1-LIKE HOMEBOX GENE (KNAT3), SWINGER (SWN), and stress recovery: COLD-REGULATED (COR413-PM1), VEGETATIVE STORAGE PROTEIN (VSP1/2) and EARLY RESPONSIVE TO DEHYDRATION (ERD3) (Figure S7B). Their coordinated reactivation in Col-0, *ein3-1*, and *atg2-1 ein3-1* suggests that this recovery module is activated independently of EIN3. Moreover, the failure to upregulate these proteins in *atg2-1* likely contributes to the impaired post-submergence recovery phenotype (Figure 6A).

Lastly, a set of proteins grouped into 2 clusters showed high abundance under non-treated conditions, followed by a mild and heterogeneous response to submergence across all genotypes, typically leaning toward slight downregulation (Figure 7). During recovery, *atg2-1* displayed a strong downregulation of these proteins when compared to the other genotypes. Notably, GO term enrichment for these clusters revealed a strong overrepresentation of chloroplast-and pigment-related processes like CHLORINA (CH1), GLUTAMATE-1-SEMIALDEHYDE 2,1-AMINOMUTASE (HEMA1), PROTOCHLOROPHYLLIDE OXIDOREDUCTASE B (PORB), GENOMES UNCOUPLED (GUN4&5), PLASTID ATPASE ASSEMBLY (PAA2). In addition, we also detected proteins functioning in redox and nitrogen metabolism: NITRATE REDUCTASE 1 (NIA1), FATTY ACID DESATURASE 5 (FAD5), BETA-AMYLASE (BAM2), auxin signaling and transport: PIN-FORMED (PIN7) ATP-BINDING CASSETTE SUBFAMILY B TRANSPORTER (ABCB19), and transcriptional or chromatin regulators like TEOSINTE BRANCHED (TCP3), and RELATIVE OF EARLY FLOWERING (REF6)(Figure S7C). As these proteins are linked to reestablishment of photosynthetic competence after submergence, their marked downregulation in *atg2-1* seems to be connected to the severe loss of chlorophyll and impaired growth seen above (Figure 6A and B, Figure S5A and B, Figure S8).

Collectively, these findings point to a dual role for autophagy: first, in dismantling sustained stress programs, particularly those governed by EIN3, and second, in enabling the reactivation of growth and metabolic functions essential for recovery. Importantly, because the proteome misregulation seen in *atg2-1* is alleviated in the *atg2-1 ein3-1* mutants, this reveals that the persistence of EIN3-dependent stress responses in *atg* mutants seems to be the primary cause for submergence and recovery susceptibility. Therefore, we propose that autophagy is required to terminate these responses once the stress has passed, allowing plants to resume growth.

Autophagy deficient plants are very sensitive to darkness so the dark submergence phenotype we observed might be influenced by caloric stress ^41^. We therefore submerged Col-0, *atg2-1*, *atg2-1 ein3-1*, *atg7-3, ein3-1* and *atg7-3 ein3-1* with full light (125 µmol m ² s ¹). As expected, all lines performed better in light submergence with much less visible senescence (Figure S8 A and B, Figure S9 A and B). In agreement with ^3^, *atg* mutants displayed reduced survival rate and mass during light submergence. Importantly, this reduction in growth was recovered in *atg2-1 ein3-1* and *atg7-3 ein3-1* (Figure S8 C and D; Figure S9 C and D), indicating that autophagy dependent regulation of EIN3 is also important for submergence responses in the light. It should be also noted that because EIN3 KO alleviates *atg*’s sensitivity to light or dark submergence, this indicates that while caloric restriction impacts plant recovery, the effect of EIN3 misregulation in *atg* mutants is the more prominent factor associated with poor survival in these mutants.

## Discussion

Given the importance of EIN3 to the execution of several developmental and stress responses, it is important to understand how cells optimize EIN3 activity and downstream responses. Our study reveals a yet uncharacterized mechanism behind EIN3 proteostasis, namely how autophagy modulates EIN3 levels (Figure 1 and 2). These results add a new layer of complexity to EIN3 regulation, which was previously shown to be degraded by the proteasome ^23,24^. While our data also confirms the participation of the proteasome in EIN3’s degradation (Figure 1), the dynamic and rapid nature of its turnover, such as in the presence of light (Figure 4A, ^21^), supports the idea that different proteostasis pathways work additively to effectively and quickly control EIN3 levels. Although our data clearly demonstrate that EIN3 associates with ATG8 and is degraded via autophagy, the precise molecular determinants of this interaction remain uncharacterized.

While the present study focuses on the physiological significance of EIN3 turnover in development and stress recovery, future work should address how recognition occurs, e.g., via predicted AIM motifs in EIN3 or through adaptor proteins as seen with other TFs (^11,15^).

Accumulation of EIN3 has previously been shown to cause early senescence similar to autophagy deficient mutants^4,5,20^. Interestingly, past studies^4^ reported that disrupting EIN2 in *atg5-1* lead to no visible suppression of the early senescence, suggesting that ET was not responsible for the early senescence of these mutants. On the contrary, our findings (Figure 5 Error! Reference source not found.) and other pieces of data as discussed by^18^ point to a direct connection among ET, senescence and autophagy. Because EIN3 is epistatic to EIN2^42^, it is likely that EIN3 still accumulates in *atg5-1* even after EIN2 is disabled, leading to early senescence. Interestingly, EIN3 has been shown to accumulate after salt treatment via EIN2-independent manner, namely due to the degradation of EBF1&2^43^. By analogy, interference with the autophagic degradation of EIN3 could bypass EIN2, thereby causing the early senescence observed in the *atg5 x ein2* double mutant. Additionally, and keeping with Yoshimoto’s^4^ and our past findings ^5^, *atg* senescence is abrogated by mutations in NONEXPRESSOR OF PATHOGENESIS-RELATED GENES 1 (NPR1), a regulator of the immunity plant hormone salicylic acid. Because NPR1 has been shown to interact and interfere with EIN3-dependent responses ^44,45^, it is possible that NPR1 mediated repression of early senescence in *atg* mutants is due to its interference with EIN3 signaling. While a connection to salicylic acid regulation cannot be ruled out, several lines of evidence suggest that ET contributes, at least in part to this phenotype. Firstly, NPR1 and EIN3 have been shown to interact and coregulate a set of genes involved in growth and stress responses^32^, and thus it is possible that ET-triggered senescence via EIN3 requires its interaction with NPR1 for initiation or maintenance. Secondly, recent work linking autophagy, ET and ripening ^46^, a process related to senescence, argues in favor of ET-induced senescence in *atg* mutants. Lastly, our data points to the fact that different ET-related phenotypes in *atg* mutants are dependent on EIN3. Collectively, it seems more likely that the early senescence of *atg* mutants is indeed caused at least in part by misregulation of EIN3-dependent ET signaling.

Another phenotype we observed was the differences in hypocotyl elongation during skoto-and photomorphogenesis in *atg2* and other genotypes (Figure 4 D, E). Hypocotyl elongation in darkness is regulated by ET, and the *ein3-1* mutant shows reduced ethylene-induced hypocotyl elongation ^19^. However, the authors observed that, without ET treatment, hypocotyl elongation in *ein3-1* remained unaffected, which is similar to our results. Those authors suggested that EIN3 close homologue ETHYLENE-INSENSITIVE3-LIKE (EIL1) may compensate for EIN3’s function. The overlap between EIN3 and EIL1 is supported by the fact that oxEIL1 complements *ein3-1* and also that a stronger ET-insensitivity phenotype is achieved in the double *ein3 eil1* as compared to either of the single mutants^47,48^. The restoration of wild-type-like elongation in the *atg2-1 ein3-1* double mutant suggests that EIN3 accumulation may be responsible for the shortened hypocotyl in *atg2-1*. However other studies have observed no differences in hypocotyl length for dark grown *atg5-3* and *atg7-2* after 7 days, yet they displayed a shorter hypocotyl under carbon starvation ^49^. This could be explained by differences in growth conditions from 3 days in our study to 7 days ^50^. It is possible that with 4 extra days of growth, the slower elongating *atg* mutants eventually catch up to the wild type whose hypocotyl elongation might already have plateaued, thus explaining the difference between studies.

Submergence stress enacts different responses coordinated by ET signaling and, despite evidence points to a regulatory role for EIN3 in this context^51–54^, surprisingly little is known about how this TF impacts submergence tolerance. Similarly, while autophagy is recognized as a multi-stress tolerance mechanism, the scarce existing literature indicates that autophagy is needed for submergence tolerance^3^. Our data indeed indicates that autophagy is necessary for full tolerance to submergence, and that this is mainly achieved through timely downregulation of EIN3-dependent responses (Figure 6, 7). Our proteome analysis revealed that autophagy deficiency leads to an EIN3-dependent accumulation of hypoxia-stress and cell wall-modification related proteins during the recovery stage (Figure S7A). The accumulation of these proteins during the submergence stage is probably required to assist with the metabolic, gas diffusion and structural changes connected to growth restriction ^51,55–58^, their persistence in the *atg2-1* mutant demonstrates an inability to terminate this stress-related programs once the stress has been removed. This observation is in line with the fact that the proteome difference between *atg2* and the other genotypes became more apparent after submergence and recovery stage, which functions as consecutive stresses which autophagy mutants do not tolerate well^1^.

Connected to growth restriction, we observed downregulation of pigment and photosynthesis related proteins during submergence which was further magnified during *atg2-1* (Figure S7C). A rapid submergence-induced downregulation of chlorophyll-related genes/proteins and photosynthesis is connected with a quiescent tolerance strategy ^59^, and it should be overturned once submergence subsides^60^. Importantly, maintaining photosynthetic assembly during submergence is instrumental for rapid reinitiation of growth and thus an efficient recovery ^61^. In opposition, leaf senescence of submerged leaves has been shown to further magnify systemic photosynthetic shutdown^62^ and this seems to be in line for what we report for the *atg* mutants undergoing submergence and recovery (Figure 6, Figure S5,8 and 9) and recent findings using *atg5-1*^63^. Importantly, ET promoted, EIN3-mediated signaling is connected to leaf senescence during re-oxygenation, via an age-dependent mechanisms which controls termination of older leaves as potential source of resources to restart growth^28^. We have previously shown that combinatorial stresses can “mimic” premature aging in *atg* mutants and lead to unrestricted death^5^, so it is likely that EIN3 accumulation and the contrasting stresses experienced under submergence and recovery coalesce to promote exaggerated senescence in these mutants (Figure 6) which prevents effective recovery. Concurrently, we also observed that *atg2* failed to accumulate cell wall-related biosynthetic genes and key developmental TF like BES1 and HY5 during the recovery stage. Given the extent of modifications cell walls undergo during submergence (e.g.^64,65^), it should be expected that during the recovery stage cell wall biogenesis is reenacted to restore cell wall integrity and resume growth. Cell wall recovery occurs through a combination of the novo synthesis, modification and salvaging of components through internalization and recycling ^66^, it is not unlikely that autophagy is required to some extent to facilitate this step and thus autophagy deficiency has a negative impact on the cells capacity to upregulate cell wall regenerative pathways. Besides structural role, cell wall also functions as a signaling hub for developmental regulation, in particular OsWAK11 has been shown to trigger adaptation to environment through monitorization of cell wall status and regulation of brassinosteroid signaling thus modulating growth responses^67^. Precisely, our proteome analysis indicates that WAK1/2, BES1 and HY5 are downregulated in *atg2* mutants during recovery, opening to speculation that defects in sensing cell wall status might contribute to the prolonged activation of submergence programs seen in this mutant and concomitant recovery delays.

Intriguingly, our comparative proteomic analysis of the *atg2* single mutant and the *atg2 ein3* double mutant revealed that GO terms associated with autophagy were consistently downregulated in the absence of EIN3, independent of treatment. This unexpected observation raises the possibility that EIN3 may play a previously unrecognized role in promoting autophagy, either through direct transcriptional regulation or via indirect effects on proteostasis and stress signaling. While this falls outside the scope of the current study, it suggests the existence of a feedback mechanism in which EIN3 not only serves as an autophagy substrate but also contributes to the expression of autophagy-related genes. Future work will be needed to determine whether EIN3 directly regulates components of the autophagy machinery, and how such a feedback loop might influence stress adaptation dynamics.

In conclusion, our findings reveal that autophagy regulates EIN3 and highlights the diverse phenotypic consequences of EIN3 misregulation in autophagy-deficient plants (Figure 8). This offers new insights into autophagy’s role as a fine-tuning mechanism for various protein programs with potential impact to the design of submergence resistant plants.

**Figure 8:**
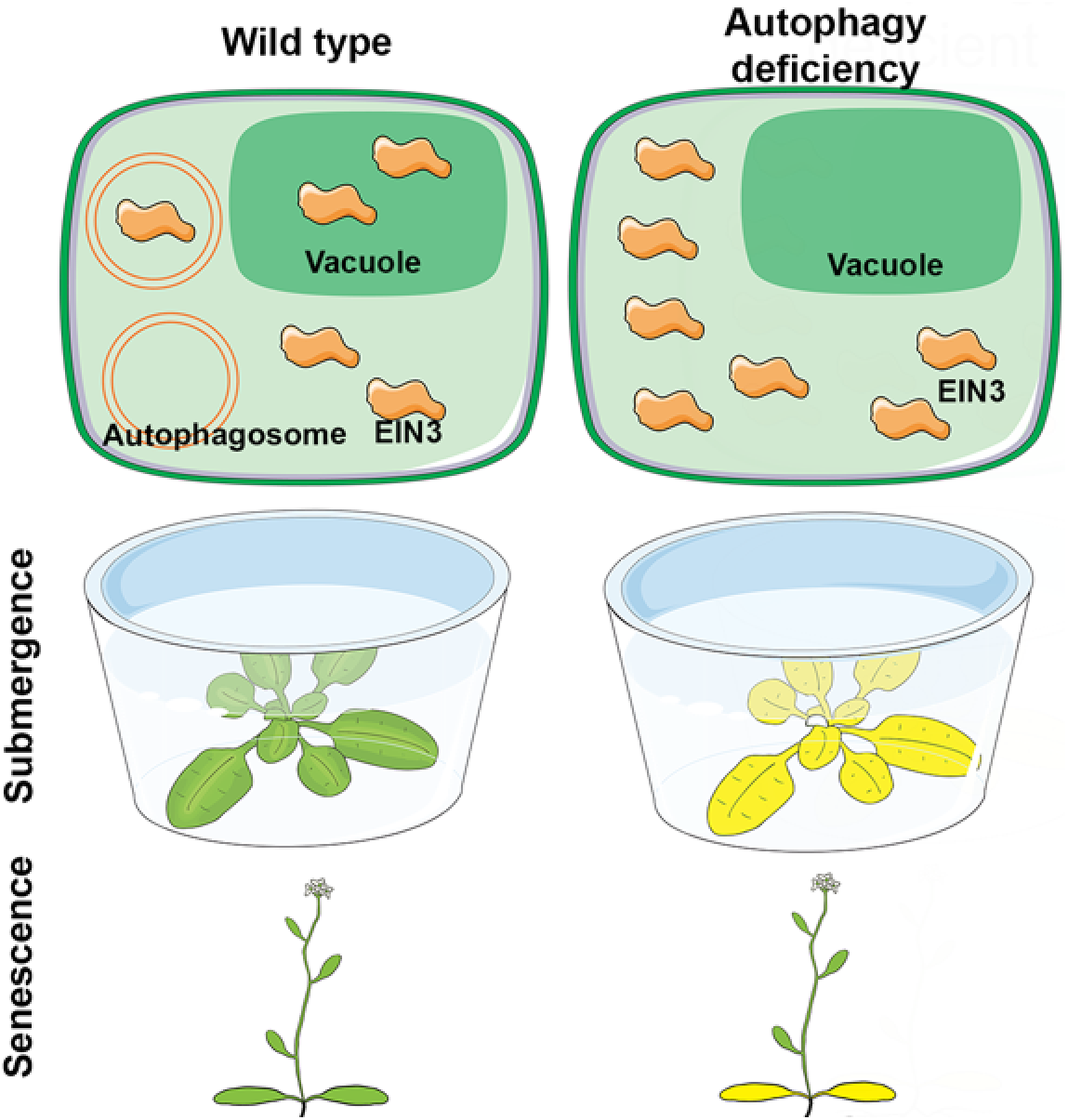
A hypothetical model displaying the regulation of EIN3 and their resulting phenotype in wild-type and autophagy deficient mutants.

## Methods

### Plant material and phenotypic characterization

Arabidopsis seedlings were grown on solid, half-strength MS media (0.8% agar, 1% sucrose, pH 5.7) and grown at 21 C° with a photoperiod of 16 h (120 μE/m^2^/s). Seeds were sterilized with chlorine gas. For experiments with 4-and 5-week-old *Arabidopsis* plants 5-day-old seedlings were moved from half-strength MS media to soil under similar conditions. The following lines were used: Col-0, *atg2-1* (SALK_076727), *atg7-3* (SAIL_11_H07), *ein3-1*^47^, *pEIN3::gEIN3-3xGFP* ^68^, *mCherry-ATG8a* ^16^. Seedlings used for colocalization and plants used for submergence were kept on half-strength MS media for 2 weeks and then placed in container and filled with tap water for one day. MG132 treatment was done as described in^11^ For ACC treatment 5-day-old seedlings were moved to liquid half-strength MS media. After 5-7 days of acclimation the media was replaced with either DMSO for control or ACC containing half-strength MS media. For dark to light the plants were grown similarly as the ACC treatment but before the lights were turned on in the growth chambers the plants were treated with E-64D (10 μM) and pepstatin A (1 μM) and immediately imaged. For the hypocotyl growth rate assay, seeds were directly put on half strength MS media and grown horizontally overnight and then rotated to be grown vertically inside of a dark box for 3 days on the SPIRO platform ^30^. After the indicated dark period, the box was opened, and the plates were exposed to standard light growth conditions. Apical hook angle determination was done as described in ^69^, in brief, seeds placed in MS or MS + 10 μM ACC, vernalized 96 h, and placed in the dark at 21°C for 4 d before pictures were taken. Apical hook angle is defined as 180° minus the angle between the tangential of the apical part with the axis of the lower part of the hypocotyl, in the case of hook exaggeration, 180° plus that angle is defined as the angle of hook curvature^70^.

For dark submergence 4-week-old plants as described above were put in plastic boxes and covered with tin foil for 24 hours with 5-10 cm room temperature tab water above the plants. At the end of the submergence the plants were taken out of the boxes and back to the tray and growth conditions they came from to let them recover.

For light submergence *Arabidopsis thaliana* Col-0, *ein3-1*, *atg2-1*, *atg7-3*, *ein3-1/atg2-1*, *ein3-1/ atg7-3* lines were sown in 5.5 x 5.5 cm pots prepared with 1:1 ratio of soil and sand and stratified in the dark for 4 days in cold room (4°C). Following stratification, plants were transferred to a growth room at 21°C and 60% humidity with a 12-12 hours day/night cycle, watered every 2-3 days, and grown for 3 weeks before the onset of the experiment. For the submergence treatment, 50 x 60 cm trays were filled with 15 liters of water a day before the experiment to allow evaporation of the dissolved oxygen in water. Before submergence, only plants that were homogenous in terms of growth were chosen for the experiment. Each tray containing an equal number (n=16) of the 4 genotypes were subjected to submergence under light for 4, 6 and 8 days in case of *atg7-3* and 8, 10 and 12 days in case of *atg2-1* or kept under light in case of control tray. A total of 4 trays were used for the experiment, 1 per submergence time-point and a control. In case of *atg2-1* group of experiments, 3 (control, 8-and 10-days sub.) trays had less than 16 plants per genotype due to lack of homogenous growth. Plants were de-submerged after respective days and allowed to recover for 5 days before phenotypic characterization. The control tray was quantified after the last time-point (8 days of submergence + 5 days of recovery).

Plants were characterized based on survival (%) and shoot fresh weight (%). For survival assay, selection criteria such as presence of meristem, signs of wilting, turgidity/ tissue degradation were followed. Survival percentage for each genotype and time-point was calculated as: Number of plants alive/Total number of plants x 100. For measuring fresh weight, plants were excised from the shoot carefully and measured on a weighing scale which was considered as absolute shoot fresh weight. Relative fresh weight percentage was calculated as: Absolute fresh weight of a genotype under a given submergence time-point/ Average of the absolute fresh weight of the genotype under control condition x 100.

### Chlorophyll extraction

Leaf samples from the same developmental stage were incubated in 80% acetone for 24 hours and then centrifuged for 5 min at 15,000g. Chlorophyll concentration was estimated following the Lichtenthaler’s equation ^71^.

### Confocal imaging

Confocal imaging of seedlings was taken with LSM-700 Zeiss confocal microscope from the Center for Advanced Bioimaging (CAB) Denmark. Images were taken using 20x air or 63x water objectives.

### Protein extraction

Proteins were extracted using 3x SDS buffer (30% glycerol, 3% SDS, 94nM Tris pH6.8, Bromophenol blue, Complete ultra-tablets EDTA-free protease inhibitor cocktail (Roche), 50 mM DTT). The samples were centrifuged for 20 min at 4 °C 14000 rpm and the supernatant was transferred to a new tube. Samples were denatured at 95 °C for 5 min before being loaded for SDS-PAGE.

### SDS-PAGE and western blotting

Protein samples were separated in a 4–20% pre-casted gel (Bio-Rad) and transferred overnight onto a PVDF membrane (GE Healthcare). Membranes were blocked for 1 hour with 5% skimmed milk (Merck) in TBST (50 mM Tris-HCL, pH 7.5, 150 nM NaCl, 0.1% Tween-20 (Sigma)) and incubated for 2 hours at room temperature with primary antibodies: anti EIN3 (R3412-2, Abiocode; 1:1000), anti-SAG12 (AS14 2771 Agrisera; 1:2000), anti-SUS1 (AS15 2830, Agrisera; 1:10000), anti ADH (AS10 685, Agrisera; 1:3000), anti-GFP (AS21 4696, Roche; 1:1000), anti-ATG8 ^72^. Membranes were washed with TBST, followed by incubation in anti-rabbit HRP conjugate or anti-mouse AP conjugate (Promega; 1:5000). For HRP detection: Homemade chemiluminescent substrate ^73^ was applied before detection. For AP detection: NBT/BCIP (Roche) was applied to the membrane until bands were visible. Bands were quantified using ImageJ and normalized to loading controls.

### Immunoprecipitation

Roughly 1g of 14-day-old seedlings were ground with mortar and pestle and 2 ml of IP extraction buffer (50 mM Tris-HCL, pH 7.5, 150 nM NaCl, 5% glycerol, 1 mM EDTA, 0.1% NP40, 10 mM DTT, Complete ultra-tablets EDTA-free protease inhibitor cocktail (Roche). The samples were centrifuged for 20 min at 4 °C 14,000 rpm and the supernatant was transferred to a new tube. The supernatant was incubated with 20 μl GFP trap beads (Chromotek) rotating for 2– 4-hours at 4 °C. The beads were washed 3 times with IP washing buffer (20 mM Tris-HCl, pH 7.5, 0.15 M NaCl, 0.1% NP40) and the washing buffer was discarded. 30 μL of 3x SDS was added to the beads and the samples were boiled for 5 min at 95 °C before being loaded for SDS-PAGE.

### LC-MS Sample preparation

Tissue samples were lysed by the addition of lysis buffer (2% SDS, 100 mM Tris pH 8.5) with a BeatBox homogeniser for 20 min at standard setting. Samples were incubated at 95 °C for 10 min and lysed with a second round of BeatBox for 10 min at standard setting followed by incubation at 95 °C for 10 min. Samples were centrifuged at 11000 xg for 10 min and transferred to a plate. Protein concentration was calculated by BCA assay (Pierce). 100 μg of each lysate were reduced with 5 mM Tcep (final conc.) for 15 min at 55 °C and alkylated with 20 mM CAA (final conc.) for 30 min at room temperature. Proteins were digested using the PAC protocol performed with the KingFisherApex robot (Thermo Fisher Scientific) in 96-well format, as described by ^74^. Briefly, the 96-well comb was placed in plate #1, samples in plate #2 in a final concentration of 70% acetonitrile (ACN) and with 25uL magnetic MagReSyn Hydroxyl beads (ReSyn Biosciences). The samples were mixed and settled for 10 min at room temperature.

Washing solutions were placed in plates #3–5 (95% ACN) and plates #6–7 (70% Ethanol). Plate #8 contained 100 μl of the digestion solution, 50 mM Triethyl amonium bicarbonate (TEAB) with Trypsin and Lys-C in 1:50 (enzyme: protein) ratio. The digestion was set to 16 hours at 37°C. Samples were acidified after digestion to a final concentration of 1% formic acid (FA).

After, samples were desalted using EasyPep peptide clean-up plates (Thermo Fisher; catalog number: A44522) following the manufacturer’s instructions. Eluates were dried by vacuum centrifugation, resuspended in buffer A* (2% ACN, 0.1 % TFA) and quantified with Tecan Lunatic. After the quantification, samples were diluted to 0.05 μg/μl and transferred to a TwincTec plate for the MS analysis.

### LC-MS

Peptides were separated on an Aurora (Gen3) 25 cm, 75 μM ID column packed with C18 beads (1.6 μm) (IonOpticks) using a Vanquish Neo (Thermo Fisher Scientific) UHPLC. Peptide separation was performed using a 60 min stepped gradient of 2-17% solvent B (0.1% formic acid in acetonitrile) for 31 min, 17-25% solvent B for 11 min, 25-35% solvent B for 6 min, using a constant flow rate of 400 nL/min. Column temperature was controlled at 50 °C. Upon elution, peptides were injected via a Nanospray Flex ion source and 30-uM steel emitter (Evosep) into a Tribrid Ascend mass spectrometer (Thermo Scientific). The Spray Voltage was set to 2200 (V). Data was acquired in data independent mode with the Orbitrap MS resolution 60.000 for full scan range 400-900m/z, AGC target was 250%, and maximum injection time set to auto. 42 DIA scans with 12 Th width and 1 Th overlap spanning a mass range of 400-900 m/z were acquired at 15,000 MS resolution, AGC target 1000%, and maximum injection time of 27 ms. HCD fragmentation normalised collision energy (NCE) was set to 30%.

### LC-MS data analysis

MS files were processed using Spectronaut version 19.5 (Biognosys) in direct DIA search mode with the default workflow. Briefly, UniProtKB UP000006548 (Taxa ID: 3702) Arabidopsis thaliana FASTA database was used. Carbaminomethylation of cysteine was set as a fixed modification, whereas the oxidation of methionine, and the acetylation of the protein N-terminus were set as variable modifications. The maximum number of missed cleavages was 2, and the minimum peptide amino acid length was 7. The false discovery rate for PSM, peptide, and protein groups was set to 0.01. All statistical analysis was performed using in-house developed python code (Proteomelit, version 1.1.0), based on the automated analysis pipeline of the Clinical Knowledge Graph ^75^. Intensity values were log2-transformed and features with less than 3 valid values in at least one group were removed. Missing values were imputed using the MinProb approach ^76^ with a width of 0.3 and downshift of 1.8. Differentially expressed features were identified by statistical unpaired t-tests and one-way ANOVA tests, permutation-based False Discovery Rate (FDR) correction for multiple hypothesis testing with FDR 0.05, Fold-change (FC) 1, s0 1, and 250 permutations ^77^. Volcano plots display all features statistically tested, with up-and down-regulated features highlighted in red and blue, respectively.

Functional enrichment analysis of significant features was performed by applying Fisher’s exact test and Benjamini-Hocheberg correction. Gene ontology and Reactome annotations were used for the enrichment testing. The analysis was performed at the protein level. Up-and down-regulated features were used as foreground and the remaining features in the dataset were considered background.

## Data availability

The mass spectrometry proteomics data will be deposited to the ProteomeXchange Consortium (http://proteomecentral.proteomexchange.org) via the PRIDE partner repository with the data set identifier

### RNA extraction

RNA was extracted as described by ^78^. In brief, TRIzol reagent (Thermo Scientific) was used to extract RNA from plant material and then cleaned with DNase (Thermo Scientific) and reverse transcribed into cDNA by Reverse Aid (Thermo Scientific). RTqPCR was done with Quantstudio 1 (Applied Biosystems) with SYBR Green master mix (Thermo Fisher) as described by the manufacturers. Actin (ACT2) was used as a control.

### Vector construction

To create pDEST-^GW^ EIN3 VYCE we did PCR (CloneAMP HIFI PCR premix, TAKARA) of previously extracted cDNA from Col-0. The coding sequence for EIN3 was used with the flanking recognition sequence for Gateway cloning (Thermofisher). The PCR products were collected from gel extraction DNA kit (T1020L New England Biolabs) and then cloned into pDONR221 (Invitrogen). The resulting product was transferred (LR Clonase, Introgen) into the pDEST-^GW^ VYCE BiFC vectors ^79^.

### BiFC transient transformation

GV3101 Agrobacterium with the desired vectors were grown overnight in LB media with their respective antibiotics. The cultures were spun at 3,000 RPM for 15 minutes, the supernatant was discarded, and the culture resuspended in infiltration media (10 mM MgCL2, 10 mM MES-K, 100 µM acetosyringone) to reach a 600 nm density of 0.1 to 0.2. The cultures were then mixed and incubated at room temperature for 2 hours, followed by leaf infiltration into 3-4 weeks old *N. benthamiana*.

### Hypocotyl measurement

Images taken with the SPIRO platform ^30^ were analyzed using ImageJ and Tracker (https://physlets.org/tracker). Images were taken every 60 minutes for 2.5 days during darkness and every 10 minutes during continuous light.

### Triple response assay

Seeds were planted on solid ½ MS media containing either 10 mM ACC or equal volume of H_2_O and wrapped in tinfoil and kept at 4°C for 4 days. The cold treated seeds were then moved to 21°C while being kept in tinfoil for 72 hours. The angles were measured using ImageJ.

## Acknowledgements

This work was funded by a Danmarks Frie Forskningsfond grant to E.R. (DFF4307-00041B). R.C. thanks the German Academic Exchange Service (DAAD) for financial support and both R.C. and S.H. are supported by the Deutsche Forschungsgemeinschaft (DFG, German Research Foundation) under Germany’s Excellence Strategy (CIBSS—EXC-2189—Project ID 390939984). We would also like to thank Suksawad Vongvisuttikun for technical assistance. Confocal microscopy imaging was performed using equipment from the Center for Advanced Bioimaging (CAB) Denmark, University of Copenhagen. Proteomic sample preparation, LC-MS measurements, and data analysis at the University of Copenhagen (UCPH), supported by the Novo Nordisk Foundation (NNF) (grant agreement number NNF19SA0059305). Thanks are due to Koki Yoshimoto (anti-ATG8a antibody) and Anna Stepanova (*ein3-1* seeds). Arabidopsis_ShallowRoots & ArabidopsisTopView icon by James-Lloyd https://www.badgrammargoodsyntax.com/ are licensed under CC0 https://creativecommons.org/publicdomain/zero/1.0. emptycell-1, glass-water & protein-23 icon by Servier https://smart.servier.com/ are licensed under CC-BY 3.0 Unported https://creativecommons.org/licenses/by/3.0.

---

**Figure S1:**
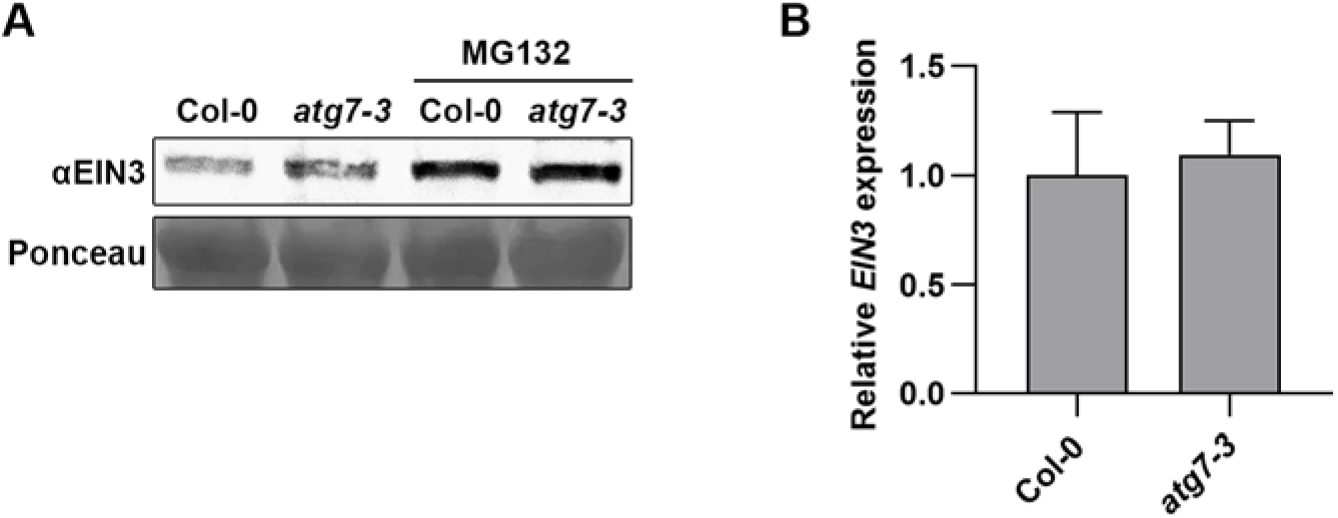
EIN3 accumulates in autophagy deficient *atg7-3* mutants. **A)** Immunoblots from protein extracts of Col-0 and *atg7-3* seedlings treated with or without MG132. **B)** Relative expression of *EIN3* in *atg7-3* and relative to Col-0. *Actin2* was used to normalize the expression. Experiments were repeated twice with similar results.

**Figure S2:**
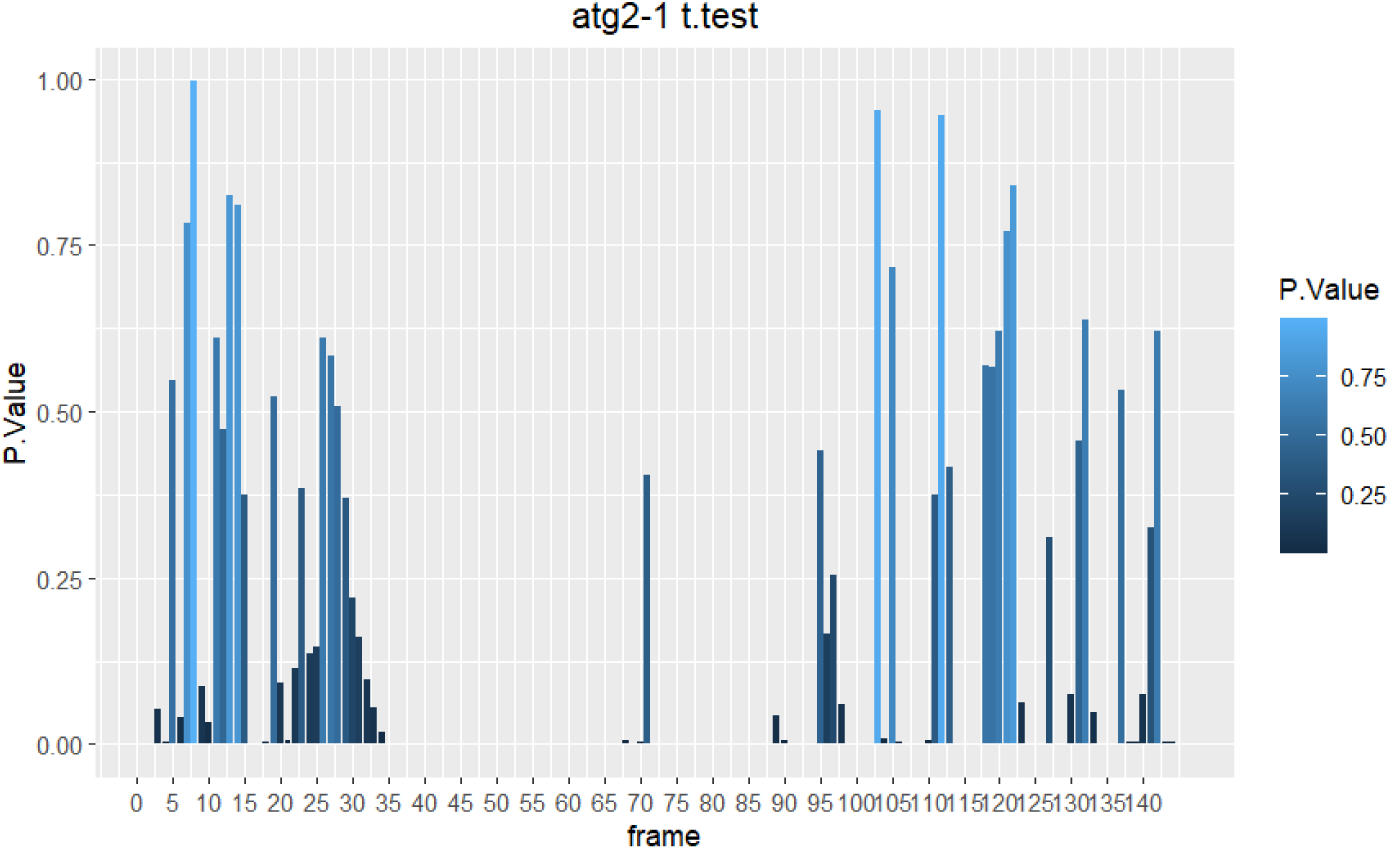
Observed differences in hypocotyl growth of Col-0, *atg2-1*, *ein3-1*, and *atg2-1 ein3-1* are not due to differential germination timing. Seedling growth was imaged every ∼5 minutes using the SPIRO automated camera system, and seedling size was quantified for each image (frame). P values presented in the figure were calculated using Welch’s t-test, comparing *atg2-1* to each other genotype at each frame.

**Figure S3:**
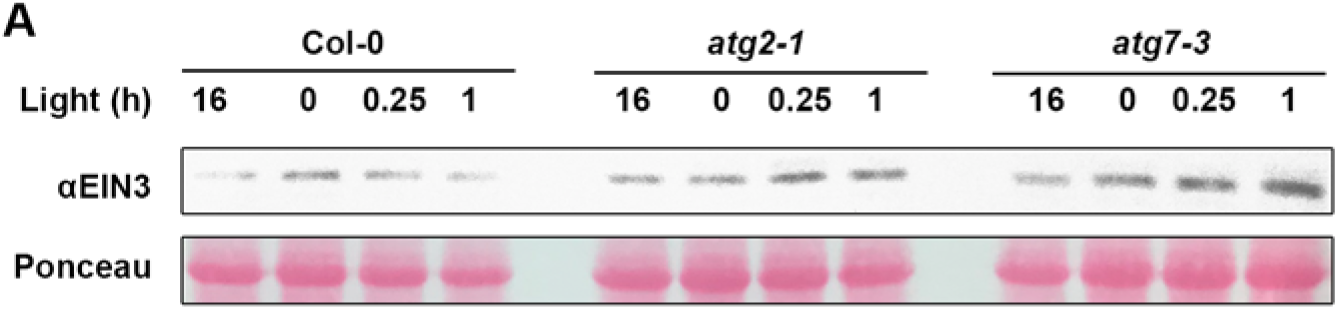
Immunoblot showing EIN3 is rapidly being degraded upon exposure to light, and this degradation is inhibited in *atg2-1 and atg7-3* mutants.

**Figure S4:**
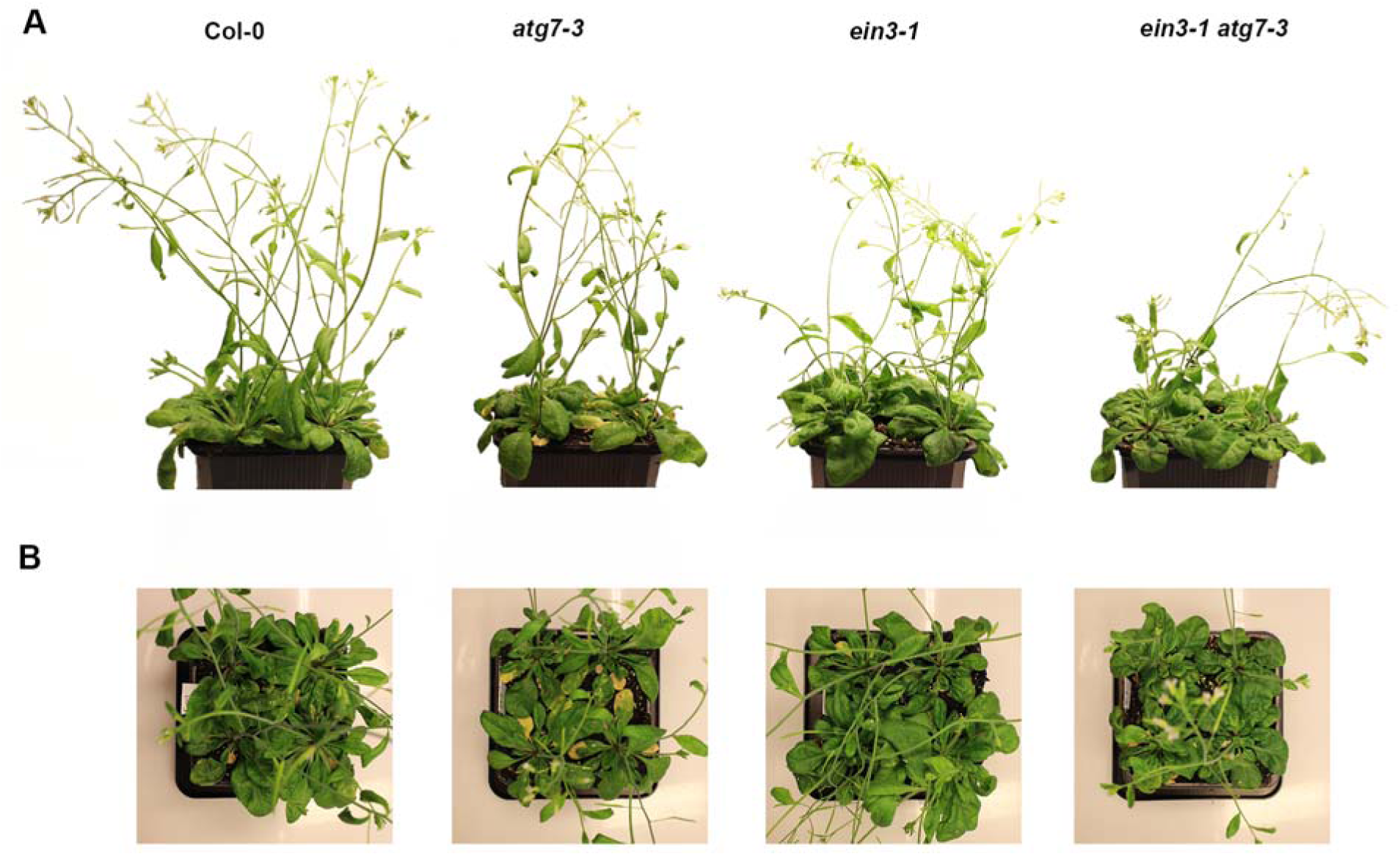
Early senescence in *atg7-3* is dependent on EIN3. **A,B)** 5-week-old *Arabidopsis* with different genotypes grown at standard conditions imaged laterally (A) or from above (B).

**Figure S5:**
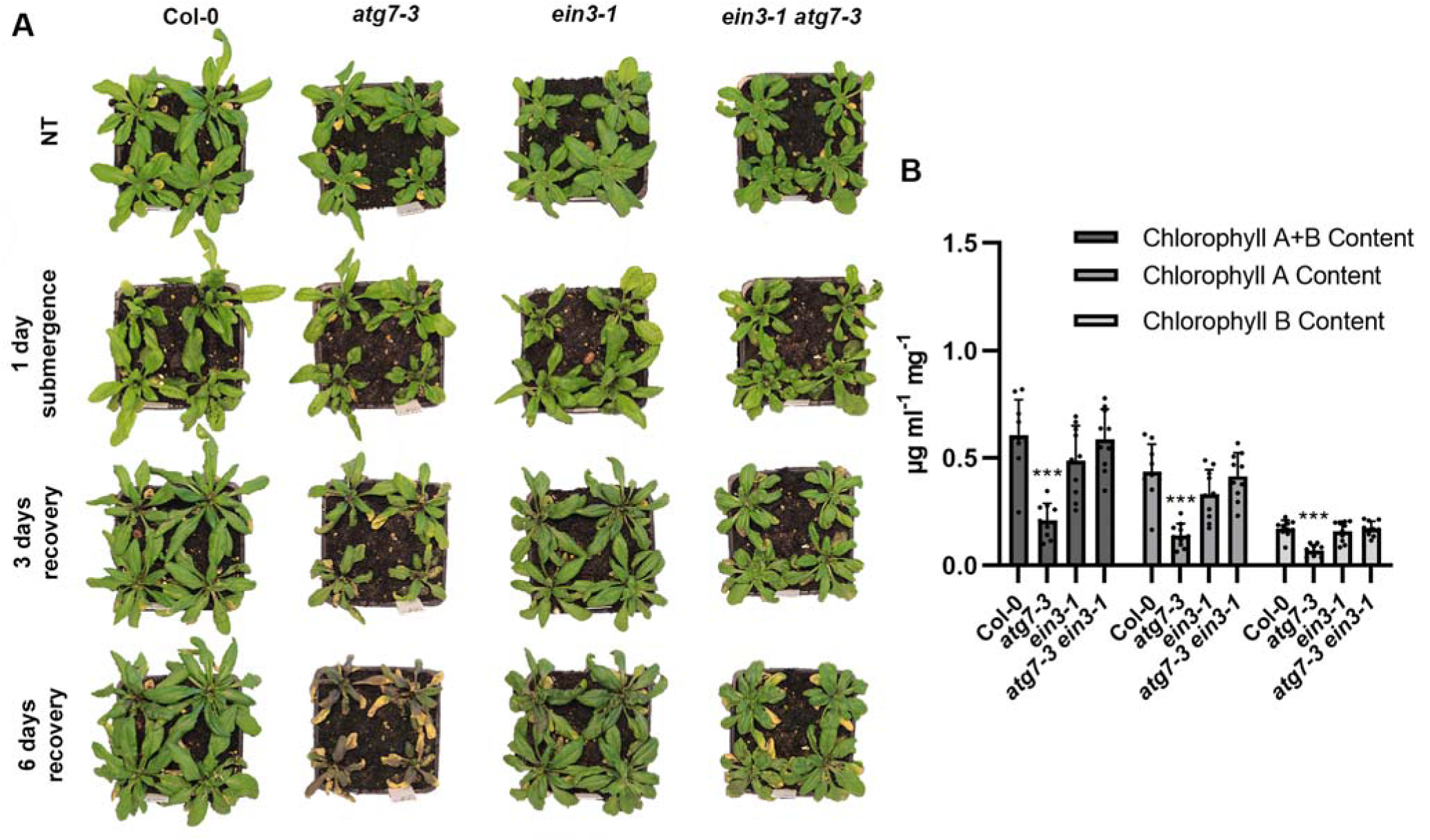
Submergence stress of *atg7-3* is EIN3 dependent. **A)** 4-week-old *Arabidopsis* before submergence, (NT), immediately after 1 day of submergence and 3-or 6-days recovery. **B)** Chlorophyll A, B, and total (A+B) content (µg mLL¹ extract mgL¹ fresh weight) after 6 days of recovery. Values are from a mean of 10 different plant rosette leaves. Asterisk depicts statistical significance from Col-0 according to a Tukey’s multiple comparison’s test (***0.001).

**Figure S6:**
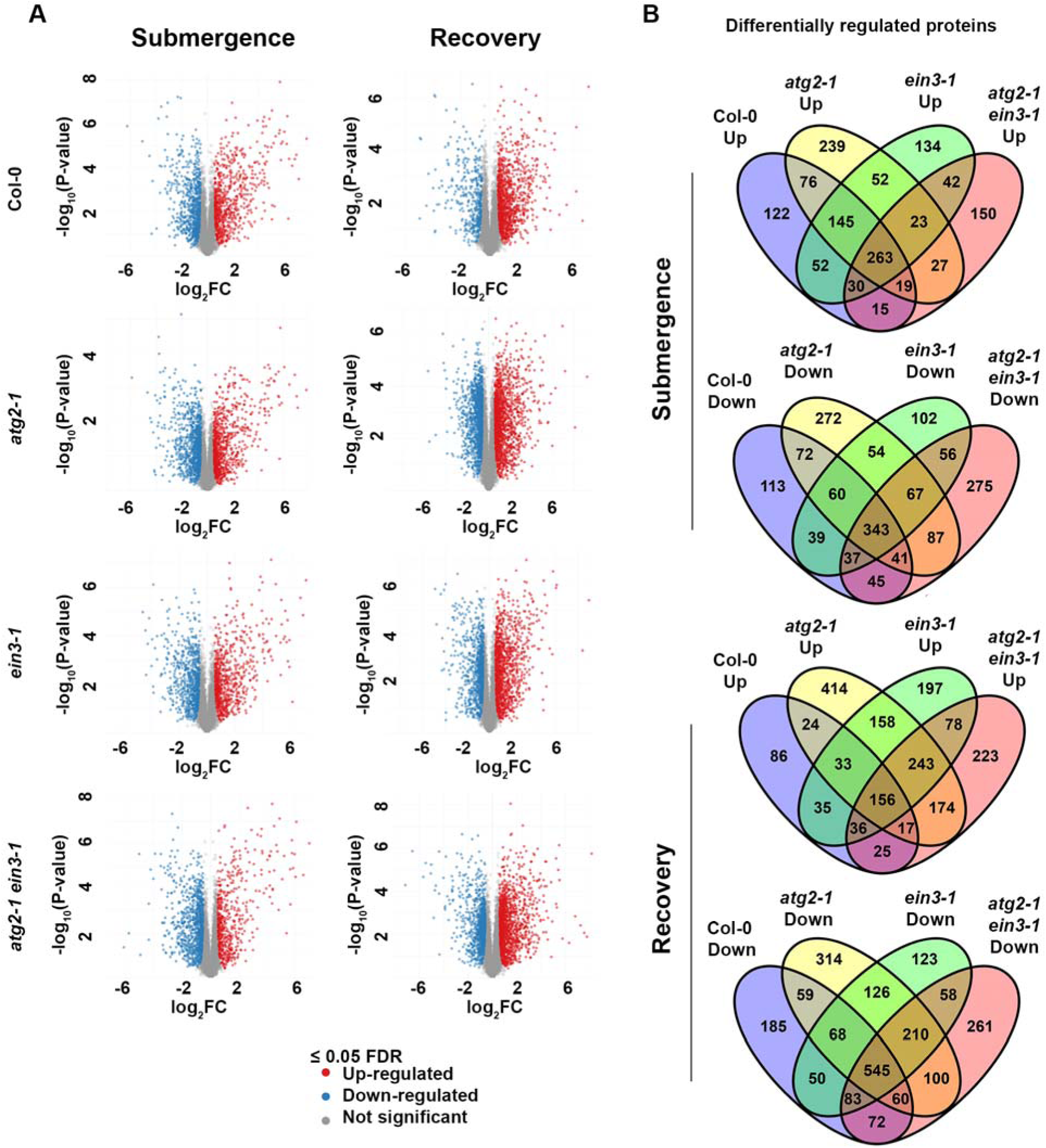
Differentially regulated proteins in Col-0, *ein3-1*, *atg2-1* and *atg2-1 ein3-1* after submergence and during recovery. **A)** Volcano plots of 4-week-old plants after dark submergence for 1 day (Submergence) compared to no treatment and volcano plots of submergence plants compared to plants which have recovered for 3 days under standard growth conditions (recovery). **B)** Venn diagram of (A).

**Figure S7:**
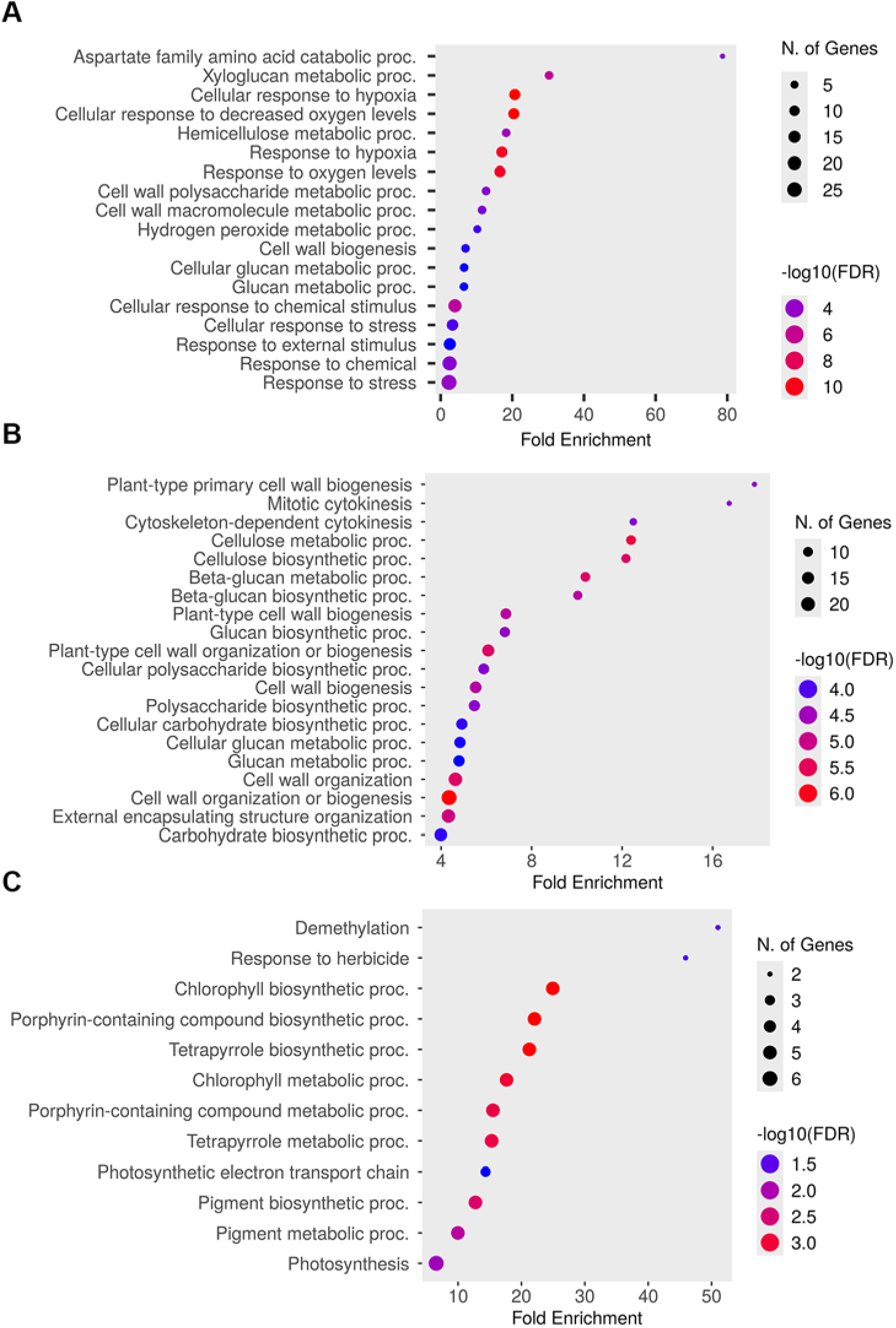
Dot plots showing top enriched Gene Ontology (GO) terms from selected protein clusters misregulated in *atg2-1* compared to all other genotypes. A) GO enrichment analysis of clusters comprising submergence-induced proteins that remain abnormally elevated in atg2-1 during recovery. B) GO enrichment of clusters containing proteins suppressed during submergence and re-accumulated during recovery in all genotypes except *atg2-1.* C) GO enrichment of clusters with proteins mildly affected by submergence but strongly downregulated during recovery specifically in *atg2-1*.

**Figure S8:**
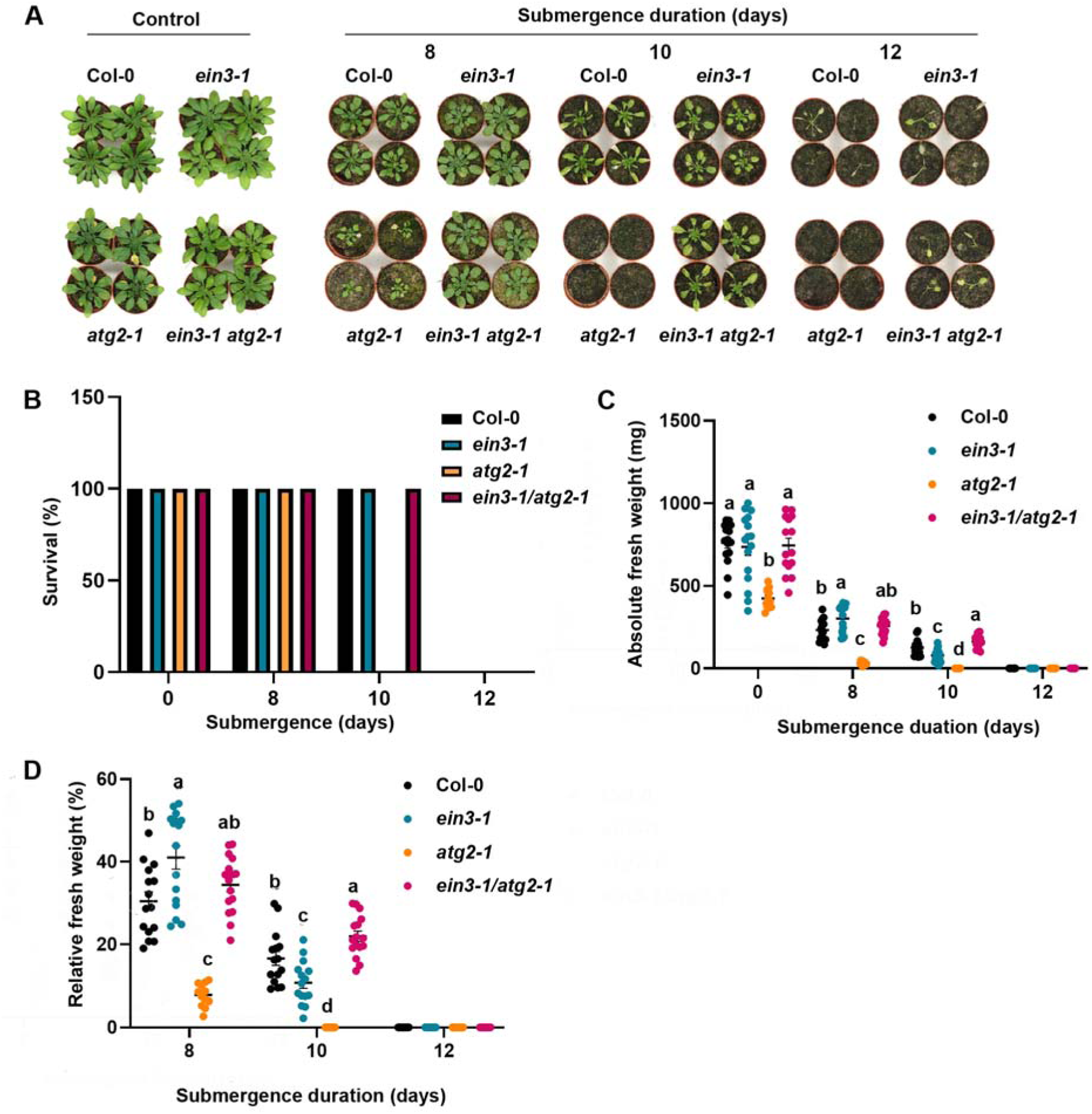
Autophagy deficiency influences submergence of *atg2-* mutants in the light. **A)** Photos of 4 weeks old *Arabidopsis* submerged for the indicated number of days. **B)** Survival rate of (A). **C)** Absolute fresh weight of (A). **D)** Relative fresh weight of (A). For C and D n=12 and results are presented as mean ± standard error of the mean (SEM), with a significance threshold set at p ≤ 0.05 as shown by compact letter display (CLD). One-way analysis of variance (one-way ANOVA) and Tukey’s multiple comparison tests were used in case of absolute and relative fresh weight data for all the genotypes under individual submergence time-points.

**Figure S9:**
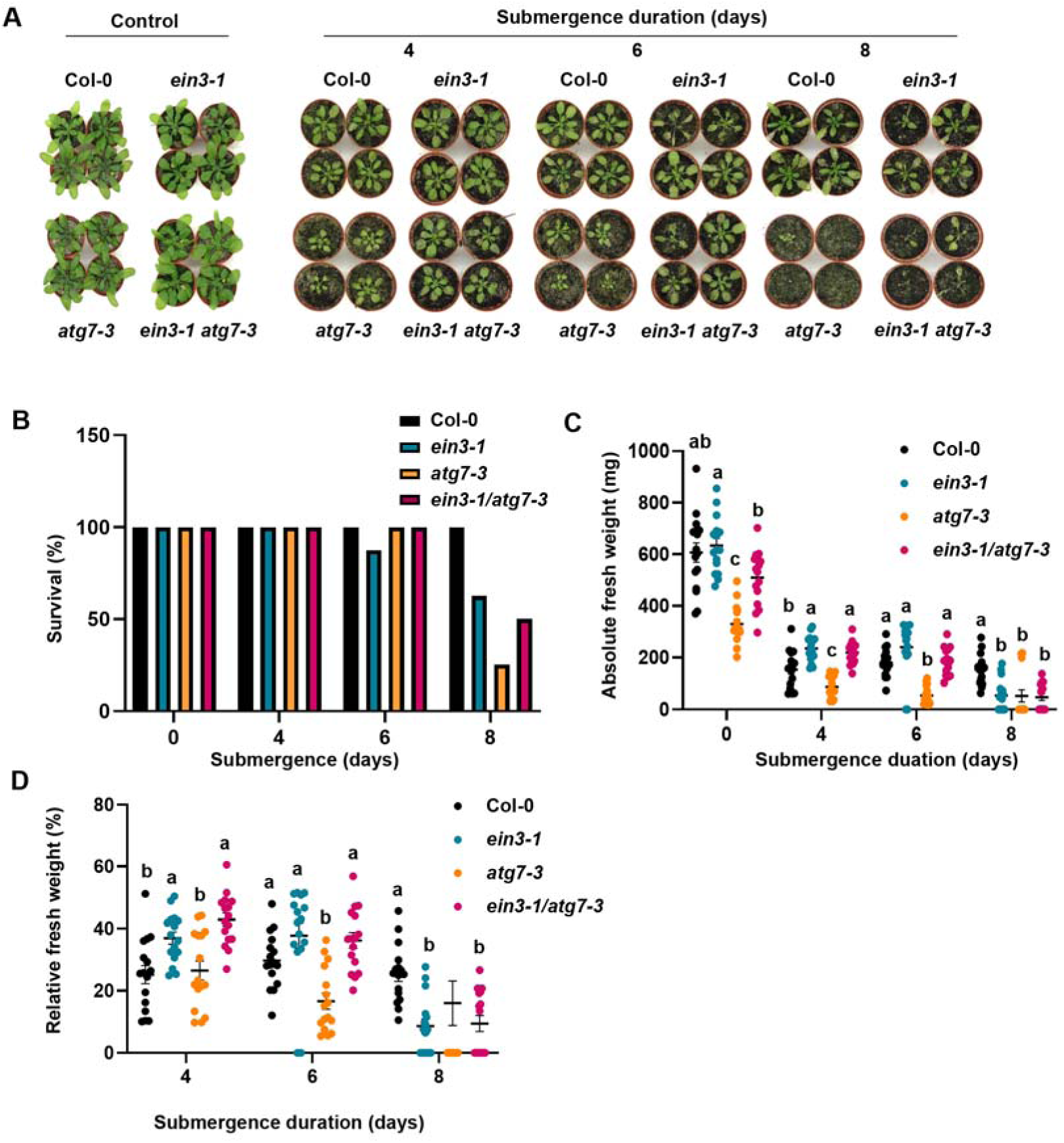
Autophagy deficiency influences submergence of *atg7-3* mutants in the light. **A)** Photos of 4 weeks old Arabidopsis submerged for the indicated number of days. **B)** Survival rate of (A). **C)** Absolute fresh weight of (A). **D)** Relative fresh weight of (A). For C and D n=16 and results are presented as mean ± standard error of the mean (SEM), with a significance threshold set at p ≤ 0.05 as shown by compact letter display (CLD). One-way analysis of variance (one-way ANOVA) and Tukey’s multiple comparison tests were used in case of absolute and relative fresh weight data for all the genotypes under individual submergence time-points.

**Table 1.**
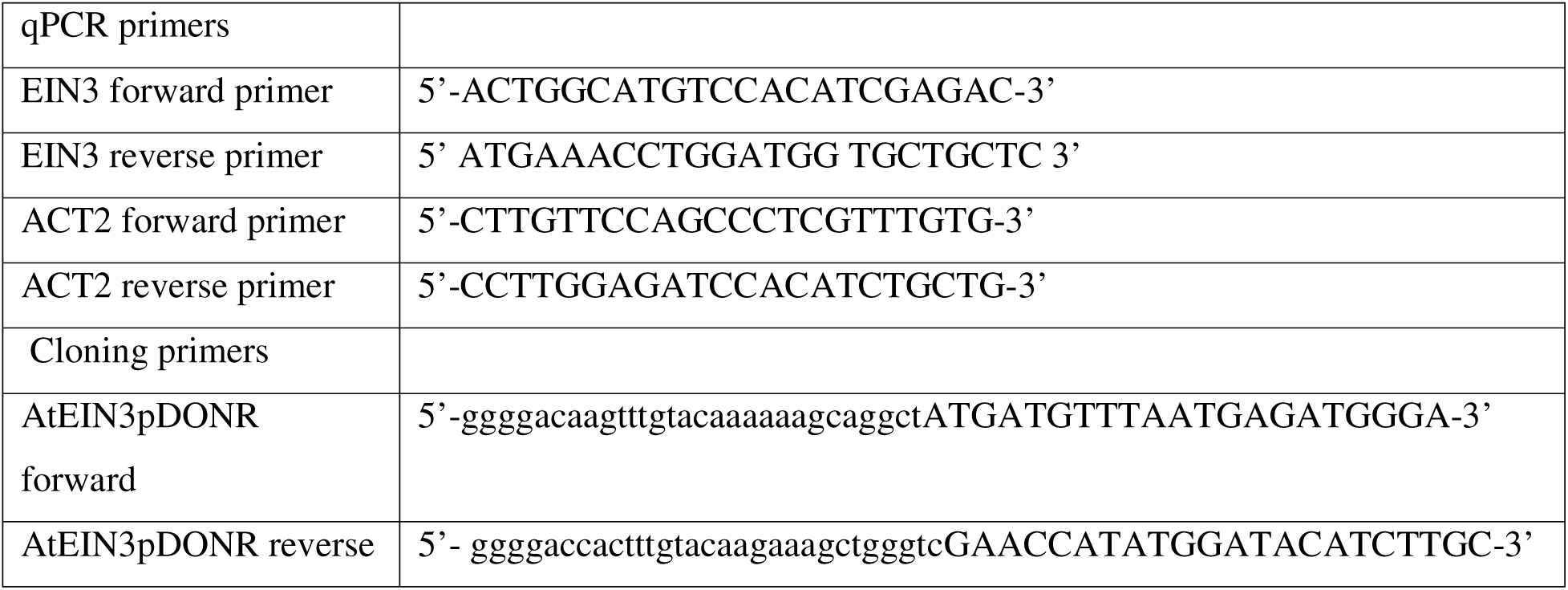
Primers used.

## References

1. Rodriguez, E. et al. Autophagy mediates temporary reprogramming and dedifferentiation in plant somatic cells. The EMBO Journal 39, e103315 (2020).

2. Yu, G. & Klionsky, D. J. Life and Death Decisions—The Many Faces of Autophagy in Cell Survival and Cell Death. Biomolecules 12, 866 (2022).

3. Chen, L. et al. Autophagy contributes to regulation of the hypoxia response during submergence in Arabidopsis thaliana. Autophagy 11, 2233–2246 (2015).

4. Yoshimoto, K. et al. Autophagy Negatively Regulates Cell Death by Controlling NPR1-Dependent Salicylic Acid Signaling during Senescence and the Innate Immune Response in Arabidopsis. The Plant Cell 21, 2914–2927 (2009).

5. Munch, D. et al. Autophagy deficiency leads to accumulation of ubiquitinated proteins, ER stress, and cell death in Arabidopsis. Autophagy 10, 1579–1587 (2014).

6. Leong, J. X. et al. A bacterial effector counteracts host autophagy by promoting degradation of an autophagy component. The EMBO Journal 41, e110352 (2022).

7. Thirumalaikumar, V. P. et al. Selective autophagy regulates heat stress memory in Arabidopsis by NBR1-mediated targeting of HSP90.1 and ROF1. Autophagy 17, 2184–2199 (2021).

8. Liu, Y., Xiong, Yan & Bassham, D. C. Autophagy is required for tolerance of drought and salt stress in plants. Autophagy 5, 954–963 (2009).

9. Yan, H. et al. Autophagy and its mediated mitochondrial quality control maintain pollen tube growth and male fertility in Arabidopsis. Autophagy 19, 768–783 (2023).

10. Yu, P. & Hua, Z. The ubiquitin-26S proteasome system and autophagy relay proteome homeostasis regulation during silique development. Plant J 111, 1324–1339 (2022).

11. Ebstrup, E. et al. NBR1-mediated selective autophagy of ARF7 modulates root branching. EMBO reports 25, 2571–2591 (2024).

12. Shinozaki, D., Takayama, E., Kawakami, N. & Yoshimoto, K. Autophagy maintains endosperm quality during seed storage to preserve germination ability in Arabidopsis. Proceedings of the National Academy of Sciences 121, e2321612121 (2024).

13. Acheampong, A. K. et al. EXO70D isoforms mediate selective autophagic degradation of type-A ARR proteins to regulate cytokinin sensitivity. Proceedings of the National Academy of Sciences 117, 27034–27043 (2020).

14. Hachez, C. et al. The Arabidopsis Abiotic Stress-Induced TSPO-Related Protein Reduces Cell-Surface Expression of the Aquaporin PIP2;7 through Protein-Protein Interactions and Autophagic Degradation. The Plant Cell 26, 4974–4990 (2014).

15. Nolan, T. M. et al. Selective Autophagy of BES1 Mediated by DSK2 Balances Plant Growth and Survival. Developmental cell 41, 33 (2017).

16. Svenning, S., Lamark, T., Krause, K. & Johansen, T. Plant NBR1 is a selective autophagy substrate and a functional hybrid of the mammalian autophagic adapters NBR1 and p62/SQSTM1. Autophagy 7, 993–1010 (2011).

17. Tarnowski, L. et al. A selective autophagy cargo receptor NBR1 modulates abscisic acid signalling in Arabidopsis thaliana. Sci Rep 10, 7778 (2020).

18. Kumaran, G. et al. Autophagy restricts tomato fruit ripening via a general role in ethylene repression. New Phytologist 246, 2392–2404 (2025).

19. Binder, B. M. et al. The Arabidopsis EIN3 Binding F-Box Proteins EBF1 and EBF2 Have Distinct but Overlapping Roles in Ethylene Signaling. The Plant Cell 19, 509–523 (2007).

20. Li, Z., Peng, J., Wen, X. & Guo, H. ETHYLENE-INSENSITIVE3 Is a Senescence-Associated Gene That Accelerates Age-Dependent Leaf Senescence by Directly Repressing miR164 Transcription in Arabidopsis[C][W]. Plant Cell 25, 3311–3328 (2013).

21. Shi, H. et al. Seedlings Transduce the Depth and Mechanical Pressure of Covering Soil Using COP1 and Ethylene to Regulate EBF1/EBF2 for Soil Emergence. Current Biology 26, 139–149 (2016).

22. Sasidharan, R. & Voesenek, L. A. C. J. Ethylene-Mediated Acclimations to Flooding Stress. Plant Physiol 169, 3–12 (2015).

23. Guo, H. & Ecker, J. R. Plant Responses to Ethylene Gas Are Mediated by SCFEBF1/EBF2-Dependent Proteolysis of EIN3 Transcription Factor. Cell 115, 667–677 (2003).

24. Potuschak, T. et al. EIN3-Dependent Regulation of Plant Ethylene Hormone Signaling by Two Arabidopsis F Box Proteins: EBF1 and EBF2. Cell 115, 679–689 (2003).

25. Yanagisawa, S., Yoo, S.-D. & Sheen, J. Differential regulation of EIN3 stability by glucose and ethylene signalling in plants. Nature 425, 521–525 (2003).

26. Wang, X. et al. Submergence stress-induced hypocotyl elongation through ethylene signaling-mediated regulation of cortical microtubules in Arabidopsis. Journal of Experimental Botany 71, 1067–1077 (2020).

27. Tsai, K.-J., Chou, S.-J. & Shih, M.-C. Ethylene plays an essential role in the recovery of Arabidopsis during post-anaerobiosis reoxygenation. Plant, Cell & Environment 37, 2391– 2405 (2014).

28. Rankenberg, T., et al. Differential leaf flooding resilience in Arabidopsis thaliana is controlled by ethylene signaling-activated and age-dependent phosphorylation of ORESARA1. Plant Comm 5, (2024).

29. Zhong, S. et al. EIN3/EIL1 cooperate with PIF1 to prevent photo-oxidation and to promote greening of Arabidopsis seedlings. Proceedings of the National Academy of Sciences of the United States of America 106, 21431 (2009).

30. Ohlsson, J. A. et al. SPIRO – the automated Petri plate imaging platform designed by biologists, for biologists. The Plant Journal 118, 584–600 (2024).

31. Eckardt, N. A. From Darkness into Light: Factors Controlling Photomorphogenesis. The Plant Cell 13, 219–221 (2001).

32. Huang, P. et al. Salicylic Acid Suppresses Apical Hook Formation via NPR1-Mediated Repression of EIN3 and EIL1 in Arabidopsis. The Plant Cell 32, 612–629 (2020).

33. Weaver, L. M., Gan, S., Quirino, B. & Amasino, R. M. A comparison of the expression patterns of several senescence-associated genes in response to stress and hormone treatment. Plant Mol Biol 37, 455–469 (1998).

34. Khan, M. I. R. et al. The significance and functions of ethylene in flooding stress tolerance in plants. Environmental and Experimental Botany 179, 104188 (2020).

35. Valko, A. et al. Adaptation to hypoxia in Drosophila melanogaster requires autophagy. Autophagy 18, 909–920.

36. Li, J. et al. Oxygen-sensitive methylation of ULK1 is required for hypoxia-induced autophagy. Nat Commun 13, 1–11 (2022).

37. Cho, H.-Y., Lu, M.-Y. J. & Shih, M.-C. The SnRK1-eIFiso4G1 signaling relay regulates the translation of specific mRNAs in Arabidopsis under submergence. New Phytologist 222, 366–381 (2019).

38. Tsuji, H., Saika, H., Tsutsumi, N., Hirai, A. & Nakazono, M. Dynamic and reversible changes in histone H3-Lys4 methylation and H3 acetylation occurring at submergence-inducible genes in rice. Plant Cell Physiol 47, 995–1003 (2006).

39. Hartman, S. et al. Ethylene-mediated nitric oxide depletion pre-adapts plants to hypoxia stress. Nature Communications 10, 4020 (2019).

40. Liu, Z. et al. Ethylene augments root hypoxia tolerance via growth cessation and reactive oxygen species amelioration. Plant Physiol 190, 1365–1383 (2022).

41. Minina, E. A. et al. Autophagy mediates caloric restriction-induced lifespan extension in Arabidopsis. Aging Cell 12, 327–329 (2013).

42. Chao, Q. et al. Activation of the Ethylene Gas Response Pathway in Arabidopsis by the Nuclear Protein ETHYLENE-INSENSITIVE3 and Related Proteins. Cell 89, 1133–1144 (1997).

43. Peng, J. et al. Salt-Induced Stabilization of EIN3/EIL1 Confers Salinity Tolerance by Deterring ROS Accumulation in Arabidopsis. PLoS Genet 10, e1004664 (2014).

44. Huang, P. et al. Salicylic Acid Suppresses Apical Hook Formation via NPR1-Mediated Repression of EIN3 and EIL1 in Arabidopsis. The Plant Cell 32, 612 (2019).

45. Yu, X., Xu, Y. & Yan, S. Salicylic acid and ethylene coordinately promote leaf senescence. Journal of Integrative Plant Biology 63, 823–827 (2021).

46. Kumaran, G. et al. Autophagy restricts tomato fruit ripening via a general role in ethylene repression. New Phytologist 246, 2392–2404 (2025).

47. Chao, Q. et al. Activation of the Ethylene Gas Response Pathway in Arabidopsis by the Nuclear Protein ETHYLENE-INSENSITIVE3 and Related Proteins. Cell 89, 1133–1144 (1997).

48. Alonso, J. M. et al. Five components of the ethylene-response pathway identified in a screen for weak ethylene-insensitive mutants in Arabidopsis. Proceedings of the National Academy of Sciences 100, 2992–2997 (2003).

49. Avin-Wittenberg, T. et al. Global Analysis of the Role of Autophagy in Cellular Metabolism and Energy Homeostasis in Arabidopsis Seedlings under Carbon Starvation. The Plant Cell 27, 306–322 (2015).

50. Rigault, M., Citerne, S., Masclaux-Daubresse, C. & Dellagi, A. Salicylic acid is a key player of Arabidopsis autophagy mutant susceptibility to the necrotrophic bacterium Dickeya dadantii. Sci Rep 11, 3624 (2021).

51. Fukao, T., Xu, K., Ronald, P. C. & Bailey-Serres, J. A variable cluster of ethylene response factor-like genes regulates metabolic and developmental acclimation responses to submergence in rice. The Plant Cell 18, 2021–2034 (2006).

52. Cho, H.-Y., Chou, M.-Y., Ho, H.-Y., Chen, W.-C. & Shih, M.-C. Ethylene modulates translation dynamics in Arabidopsis under submergence via GCN2 and EIN2. Science Advances 8, eabm7863 (2022).

53. Hong, C.-P., Wang, M.-C. & Yang, C.-Y. NADPH Oxidase RbohD and Ethylene Signaling are Involved in Modulating Seedling Growth and Survival Under Submergence Stress. Plants 9, 471 (2020).

54. Xie, L.-J. et al. Unsaturation of Very-Long-Chain Ceramides Protects Plant from Hypoxia-Induced Damages by Modulating Ethylene Signaling in Arabidopsis. PLOS Genetics 11, e1005143 (2015).

55. Komatsu, S., Kobayashi, Y., Nishizawa, K., Nanjo, Y. & Furukawa, K. Comparative proteomics analysis of differentially expressed proteins in soybean cell wall during flooding stress. Amino Acids 39, 1435–1449 (2010).

56. Komatsu, S. & Yanagawa, Y. Cell wall proteomics of crops. Front. Plant Sci. 4, (2013).

57. Kong, F.-J., Oyanagi, A. & Komatsu, S. Cell wall proteome of wheat roots under flooding stress using gel-based and LC MS/MS-based proteomics approaches. Biochimica et Biophysica Acta (BBA) - Proteins and Proteomics 1804, 124–136 (2010).

58. Mühlenbock, P., Plaszczyca, M., Plaszczyca, M., Mellerowicz, E. & Karpinski, S. Lysigenous Aerenchyma Formation in Arabidopsis Is Controlled by LESION SIMULATING DISEASE1. Plant Cell 19, 3819–3830 (2007).

59. Sone, C., Ito, O. & Sakagami, J.-I. Characterizing Submergence Survival Strategy in Rice Via Chlorophyll Fluorescence. Journal of Agronomy and Crop Science 198, 152–160 (2012).

60. Yeung, E. et al. A stress recovery signaling network for enhanced flooding tolerance in Arabidopsis thaliana. Proc Natl Acad Sci U S A 115, E6085–E6094 (2018).

61. Yuan, L.-B. et al. Multi-stress resilience in plants recovering from submergence. Plant Biotechnology Journal 21, 466–481 (2023).

62. Gan, L., Han, L., Yin, S. & Jiang, Y. Chlorophyll Metabolism and Gene Expression in Response to Submergence Stress and Subsequent Recovery in Perennial Ryegrass Accessions Differing in Growth Habits. Journal of Plant Physiology 251, 153195 (2020).

63. Yang, M. et al. Autophagy Regulates Plant Tolerance to Submergence by Modulating Photosynthesis. Plant, Cell & Environment 48, 2267–2284 (2025).

64. Vitorino, P. G. et al. Flooding tolerance and cell wall alterations in maize mesocotyl during hypoxia. Pesquisa Agropecuaria Brasileira 36, 1027–1035 (2001).

65. Khan, M. N. et al. Proteomic insight into soybean response to flooding stress reveals changes in energy metabolism and cell wall modifications. PLoS One 17, e0264453 (2022).

66. Barnes, W. J. & Anderson, C. T. Release, Recycle, Rebuild: Cell-Wall Remodeling, Autodegradation, and Sugar Salvage for New Wall Biosynthesis during Plant Development. Molecular Plant 11, 31–46 (2018).

67. Yue, Z.-L. et al. The receptor kinase OsWAK11 monitors cell wall pectin changes to fine-tune brassinosteroid signaling and regulate cell elongation in rice. Curr Biol 32, 2454–2466.e7 (2022).

68. Vaseva, I. I. et al. The plant hormone ethylene restricts Arabidopsis growth via the epidermis. Proceedings of the National Academy of Sciences 115, E4130–E4139 (2018).

69. Zuo, Z. et al. The mRNA decapping machinery targets LBD3/ASL9 to mediate apical hook and lateral root development. Life Science Alliance 6, (2023).

70. Vandenbussche, F. et al. The auxin influx carriers AUX1 and LAX3 are involved in auxin-ethylene interactions during apical hook development in Arabidopsis thaliana seedlings. Development 137, 597–606 (2010).

71. Lichtenhaler, H. K. & Wellburn, A. R. Determinations of total carotenoids and chlorophylls a and b of leaf extracts in different solvents. Biochemical Society Transactions 11, 591–592 (1983).

72. Yoshimoto, K. et al. Processing of ATG8s, Ubiquitin-Like Proteins, and Their Deconjugation by ATG4s Are Essential for Plant Autophagy. The Plant Cell 16, 2967–2983 (2004).

73. Mruk, D. D. & Cheng, C. Y. Enhanced chemiluminescence (ECL) for routine immunoblotting: An inexpensive alternative to commercially available kits. Spermatogenesis 1, 121–122 (2011).

74. Batth, T. S. et al. Protein Aggregation Capture on Microparticles Enables Multipurpose Proteomics Sample Preparation. Mol Cell Proteomics 18, 1027–1035 (2019).

75. Santos, A. et al. A knowledge graph to interpret clinical proteomics data. Nat Biotechnol 40, 692–702 (2022).

76. Lazar, C., Gatto, L., Ferro, M., Bruley, C. & Burger, T. Accounting for the Multiple Natures of Missing Values in Label-Free Quantitative Proteomics Data Sets to Compare Imputation Strategies. J Proteome Res 15, 1116–1125 (2016).

77. Tyanova, S. & Cox, J. Perseus: A Bioinformatics Platform for Integrative Analysis of Proteomics Data in Cancer Research. Methods Mol Biol 1711, 133–148 (2018).

78. Zuo, Z., Roux, M., Rodriguez, E. & Petersen, M. mRNA Decapping Factors LSM1 and PAT Paralogs Are Involved in Turnip Mosaic Virus Viral Infection. MPMI 35, 125–130 (2022).

79. Gehl, C., Waadt, R., Kudla, J., Mendel, R.-R. & Hänsch, R. New GATEWAY vectors for High Throughput Analyses of Protein–Protein Interactions by Bimolecular Fluorescence Complementation. Molecular Plant 2, 1051–1058 (2009).

